# Genome-wide Analysis of Bromodomain Gene Family in Arabidopsis and Rice

**DOI:** 10.1101/2022.02.08.479518

**Authors:** T.V. Abiraami, Ravi Prakash Sanyal, Hari Sharan Misra, Ajay Saini

## Abstract

The bromodomain containing proteins (Brd-proteins) belongs to family of ‘epigenetic mark readers’, integral to epigenetic regulation. The Brd-members contain a conserved ‘bromodomain’ (BRD/BRD-fold: interacts with acetylated-lysine in histones), and several additional domains, making them structurally/functionally diverse. Like animals, plants also contain multiple Brd-homologs, however the extent of their diversity and impact of molecular events (genomic duplications, alternative splicing, AS) therein, is relatively less explored. The present genome-wide analysis of *Brd*-families of *Arabidopsis thaliana* and *Oryza sativa* showed extensive diversity in structure of genes/proteins, regulatory elements, expression pattern, domains/motifs, and the bromodomain (length/sequence, location) among Brd-members. Orthology analysis identified 13 ortholog groups, three paralog groups and four singleton members. While more than 40% *Brd*-genes were affected by genomic duplication events in both plants, AS-events affected 60% *A*. *thaliana* and 40% *O*. *sativa* genes. These molecular events affected various regions (promoters, untranslated regions, exons) of different *Brd*-members with potential impact on expression/structure-function characteristics. RNA-Seq data analysis of *Brd*-members, and RT-qPCR of duplicate *Brd*-genes showed differences in tissue-specificity and response to salinity. Phylogenetic analysis based on conserved BRD-region placed the *A*. *thaliana* and *O*. *sativa* homologs into clusters/sub-clusters, consistent with ortholog/paralog groups. The bromodomain-region displayed conserved signatures in key BRD-fold elements (α-helices, loops), and variations (1-20 sites) including indels among the Brd-duplicates. Homology modeling and superposition identified structural variations in BRD-folds of divergent and duplicate Brd-members, which might affect their interaction with chromatin and associated functions. The study also showed contribution of various duplication events in *Brd*-family expansion among diverse plants, including monocots and dicots.

## 1 Introduction

Gene expression in higher animals and plants is highly complex and regulated at multiple levels in response to cellular/physiological requirements, unfavorable environmental conditions (Floris et al. 2009; Haak et al. 2017; Merchante et al. 2017; Withers and Dong, 2017). The transcription status of genes is influenced by both genetic and epigenetic components (Singh, 1998), and unlike the genetic-elements (core promoter, *cis*-elements, enhancers, silencers), the epigenetic controls involve non-sequence-based modifications to alter the expression of genes (Gibney and Nolan, 2010; Lämke and Bäurle, 2017; Rendina González et al. 2018). Epigenetic modifications of DNA and/or histones affect the state of chromatin and transcription activity (Iwasaki and Paszkowski, 2014), leading to the enhanced response potential of the genetic material (Strahl and Allis, 2000; Loidl, 2004). Post-translational modifications (PTMs: acetylation, methylation, phosphorylation, ubiquitylation etc.) affect the characteristics of several cellular proteins including histones, where such modifications modulate the nucleosome dynamics (Bowman and Poirier, 2015). Among these, acetylation of lysine plays important roles in protein-protein interactions, nuclear transport as well as modulation of chromatin state due to its effect on positive charge and steric bulk of a nucleosome (Bowman and Poirier, 2015). The cellular epigenetic regulation is based on a system of ‘writers’, ‘readers’ and ‘erasers’ proteins for dynamic management of PTM marks (Musselman et al. 2012; Zhao et al. 2018). For acetylation of lysine, lysine acetyltransferases (KATs) and lysine deacetylases/histone deacetylases (KDAC/HDACs) perform ‘writer’ and ‘eraser’ functions (Musselman et al. 2012), while the conserved bromodomain (BRD), important component of several chromatin-associated proteins, serve as ‘reader’ of lysine acetylation on histones (Drazic et al. 2016).

The bromodomain containing genes (*Brd*-genes) were first identified in the *Drosophila melanogaster* (Tamkun, et al. 1992), and subsequently reported in diverse eukaryotes (Rao et al. 2014). The ~110 amino acid bromodomain region folds into four α-helices (αZ, αA, αB, αC) and three inter-connected loops (ZA, AB, BC) to form a conserved bromodomain (BRD)/ BRD-fold with a hydrophobic pocket to recognize acetylated lysine residues in the C-terminal tail of the histones (Bottomley et al. 2004; Ferri et al. 2016; Marmorstein and Berger, 2001; Mujtaba et al. 2007). A single BRD is capable of recognizing acetylated lysine residues on different histones (Josling et al. 2012). Brd-containing proteins (alone or as multi-protein complexes) are involved in regulation of gene expression by different mechanisms viz. chromatin remodeling, histone modifications, transcriptional machinery regulation (Fujisawa et al. 2017). The *Brd*-gene family, which is a large and diverse family among different organisms, contains a total of 46 Brd-members in humans, divided into eight structurally and functionally distinct groups (Filippakopoulos et al. 2012). Human Brd-members have been studied well for their involvement in chromatin dynamics and diverse cellular functions, and have gained attention as promising drug targets for different disease conditions (Boyson et al. 2021; Cochran et al. 2019; Sanchez and Zhou, 2009; Taniguchi, 2016; Uppal et al. 2019; Zeng and Zhou, 2002).

Epigenetic-changes in chromatin structure and organization, and its impact on expression of genes is equally important for cellular and physiological requirements, and stress responses in plants (Bhadouriya et al. 2021; Fransz and De Jong, 2002; Pei et al. 2021; Rosa et al. 2013, Vergara et al. 2017). Like animals, plants also harbor a large *Brd*-gene family, however the extent of divergence of Brd-members, and their functional significance is relatively less explored. Studies on few Brd-homologs from Arabidopsis and other plants have shown their involvement in seed germination (Duque and Chua, 2003), leaf development (Chua et al. 2005), mitotic cell cycle (Airoldi et al. 2010), sugar and abscisic acid responses (Misra et al. 2017), growth and development (Kalantidis et al. 2007; Martel et al. 2017; Rao et al. 2014), pathogen perception (Sukarta et al. 2020), and as important subunits of SWI/SNF chromatic remodelers (Jarończyk et al., 2021).

The number of *Brd*-family members vary in different organisms (Rao et al. 2014). An important feature of most plants genomes is genomic duplication events that have contributed towards generation of additional copies of several genes, which can diverge towards regulatory, structural and functional difference (Barker et al. 2012; Flagel and Wendel, 2009; Qiao et al. 2019). In addition to the diversity of genes in multi-member families, it is also important to identify the conserved orthologs as well as species-specific paralogs to get insights into the evolutionary trend of genes and lineage-specific events (Altenhoff et al., 2019). Further, a large number of reports have shown, the role of alternative splicing (AS) in regulation of expression of genes in diverse conditions, and to alter key protein features among alternative isoforms (Reddy et al. 2013). Involvement of both these mechanisms on diversity of *Brd*-genes among plants is not explored well, and is worth investigating.

In the present study, we carried out genome-wide analysis of *Brd*-family in model plants *A. thaliana* (dicot) and *O. sativa* (monocot), to understand the extent of diversity of genes and proteins, regulatory regions, expression dynamics, domains-motifs architecture, and variations in the BRD-fold itself. Analysis of conserved orthologs, species-specific paralogs and singleton *Brd*-members, revealed the differential evolutionary trend of Brd-members in the two species. Result also showed the effect of genome duplication and AS-events on the characteristics of Brd genes, proteins as well as the bromodomain-fold, in both the species. Moreover, analysis also showed substantial contribution of various duplication events in *Brd*-gene copy number increase among lower and higher plants, including monocot and dicot species. To our knowledge, this is the first study on analysis of diversity of *Brd*-homologs of model plants, *A. thaliana* and *O. sativa*, which will be useful for further studies on deciphering their significance in chromatin dynamics and functions in different conditions including stress responses.

## 2 Materials and Methods

### 2.1 Identification of bromodomain genes in *A. thaliana* and *O. sativa* genome databases

Multiple databases were used for retrieval of sequence data of *Brd*-genes from *A. thaliana* (referred to as ‘*AtBrd*’) and *O. sativa* (referred to as ‘*OsBrd*’). The Arabidopsis Information Resource (TAIR, https://www.arabidopsis.org/index.jsp) and PLAZA dicots (https://bioinformatics.psb.ugent.be/plaza/versions/plaza_v4_5_dicots/, Van Bel et al. 2018) databases were used for *A. thaliana*, for *O. sativa*, Rice Genome Annotation Project (RGAP, http://rice.uga.edu/) and PLAZA monocots (version 4.5, https://bioinformatics.psb.ugent.be/plaza/versions/plaza_v4_5_monocost/, Van Bel et al. 2018) databases were used. Analysis based on Brd-family IDs (SCOP database ID: Brd superfamily, 3001843; bromodomain family, 4000871) was used for identification of Brd-family members, and confirmation was also done for presence of bromodomain at Conserved Domain Database (CDD, NCBI, https://www.ncbi.nlm.nih.gov/cdd/).

### 2.2 *In silico* analysis of characteristics of genes, proteins and transcripts

The structure and organization of *AtBrd* and *OsBrd* genes in terms of untranslated regions (UTRs), exons, and introns was analyzed using the Gene Structure Display Server (GSDS, http://gsds.cbi.pku.edu.cn/, Hu et al. 2014). The important characteristics (molecular weight, MW; isoelectric point, pI etc.) of the AtBRD and OsBRD proteins were estimated using the PROTPARAM tool on the ExPASy website (http://web.expasy.org/protparam/). The alternative isoforms of the *AtBrd* and *OsBrd* genes were retrieved from the respective databases, and compared with the corresponding constitutive isoforms by pair-wise alignment using ClustalX (Thompson et al. 1997).

### 2.3 Identification of orthologs and paralogs

For the identification of orthologs and paralogs, the Arabidopsis and rice Brd-family members by OrthoVenn2 online tool (https://orthovenn2.bioinfotoolkits.net/home, Xu et al., 2019). In brief, the BRD-protein sequences were analysed OrthoVenn2 portal (E-value, 1e-2; inflation value, 1.5) to identify the shared ortholog groups (representation of both species), species-specific paralog groups (representation of one species) and singleton sequences (not part of ortholog/paralog groups).

### 2.4 *In silico* analysis of promoter structure and *cis-*regulatory elements

Upstream (up to 2000 bp) regulatory regions of *AtBrd* and *OsBrd* genes were retrieved from the TAIR, RGAP and PLAZA databases. Presence and organization of CpG islands, transcription factor binding sites (TFBS), and tandem repeats motifs was analyzed at Plant Promoter Analysis Navigator online resource (PlantPAN3.0, http://plantpan3.itps.ncku.edu.tw/), whereas *cis-*elements (types, location, copy number) were analyzed at Plant *Cis*-Acting Regulatory Elements databases (PlantCARE, http://bioinformatics.psb.ugent.be/webtools/plantcare/html/).

### 2.5 *In silico* analysis of conserved domains, functional sites and motifs

Presence of conserved domains, important functional sites in the BRD proteins was analysed at CDD-NCBI), and PROSITE (https://prosite.expasy.org/, Sigrist et al. 2012). Domain analysis was carried out using default search parameters and only significant hits were considered for analysis. Conserved motifs were analysed at MEME online Suite (version 5.4.1, http://meme-suite.org/tools/meme, Timothy et al. 2015) using following parameters, minimum and maximum motif width: 6-50, number of motifs: 15.

### 2.6 *In silico* analysis of gene expression using RNA-Seq data

The RNA-Seq expression data (as FPKM values, Fragments Per Kilobase of transcript, per Million mapped reads) of respective *Brd*-genes was retrieved from the Arabidopsis RNA-seq Database (V2, http://ipf.sustech.edu.cn/pub/athrna/) and Rice Expression Database (http://expression.ic4r.org/). The gene/locus names were used for search and retrieval of FPKM data. For tissue-specific analysis in Arabidopsis, data was retrieved for shoot, root, stem, meristem, seedling, embryo, leaf, silique, silique, endosperm, flower and pollen. In rice, data was retrieved for root, shoot, panicle, anther, pistil, aleurone and seed. In case of multiple libraries, the average FPKM values were used. FPKM values were log2-transformed and used for generation of heat map-based transcript profiles by Heatmap Illustrator software (HemI, version 1.0, Deng et al. 2014).

### 2.7 Plant growth conditions, total RNA isolation, cDNA synthesis and RT-qPCR analysis

Seeds of *A. thaliana* ecotype Columbia-0 (Col-0) were grown on MS-agar plates containing germination media (HiMedia, India), in Sanyo MLR-351H plant growth chamber (temperature: 23 ± 1°C, a 14 h light/ 10 h dark photoperiod). For salt stress treatment, 15-day old seedlings were transferred to MS-media containing 150 mM sodium chloride, NaCl). Seeds of rice genotype NSICRc106 (obtained from International Rice Research Institute, Philippines) were grown in Hoagland media (Himedia, India) in Sanyo MLR-351H plant growth chamber as detailed previously (Sanyal et al., 2018). Six-day-old seedlings were subjected to salt stress (150 mM NaCl). Tissue samples of both the plants were collected at 24 h time-point, frozen in liquid nitrogen, and stored at −70°C. Three-five seedlings were pooled for total RNA isolation by TRIzol (Invitrogen, USA), which was assessed for quality and quantity, and treated with DNase I (Roche Diagnostics, Germany) to remove DNA contamination. Total RNA (10 μg) was reverse transcribed using SuperScript II reverse transcriptase (Invitrogen, USA) using anchored oligo(dT)35 and random nonamers (New England Biolabs, USA), as per the protocol recommended by the manufacturer.

Transcript levels of *Brd*-genes (four *AtBrd-*pairs and five *OsBrd*-pairs) were analyzed by RT-qPCR analysis using oligonucleotide primers designed utilizing the exon-intron information available from RGAP and TAIR databases (Supplementary Table 1). Briefly, RT-qPCR assays were carried out on LightCycler LC480 II (Roche Diagnostics, Germany) using SYBR Green Jumpstart *Taq* Ready mix (Sigma-Aldrich, USA) using following cycling settings: 94 °C (2 min), 45 cycles of 94 °C (15 sec), 60 °C (15 sec), 68 °C (20 sec), followed by melting curve analysis to assess the amplicon specificity. Three independent replicate sets were used and analysis was carried out as per Schmittgen and Livak (2008) using *AtActin* and *OselF1α* as reference genes. Statistical analysis was carried out by Student’s t-test and differences were considered significant only when the P < 0.05.

**Table 1.**
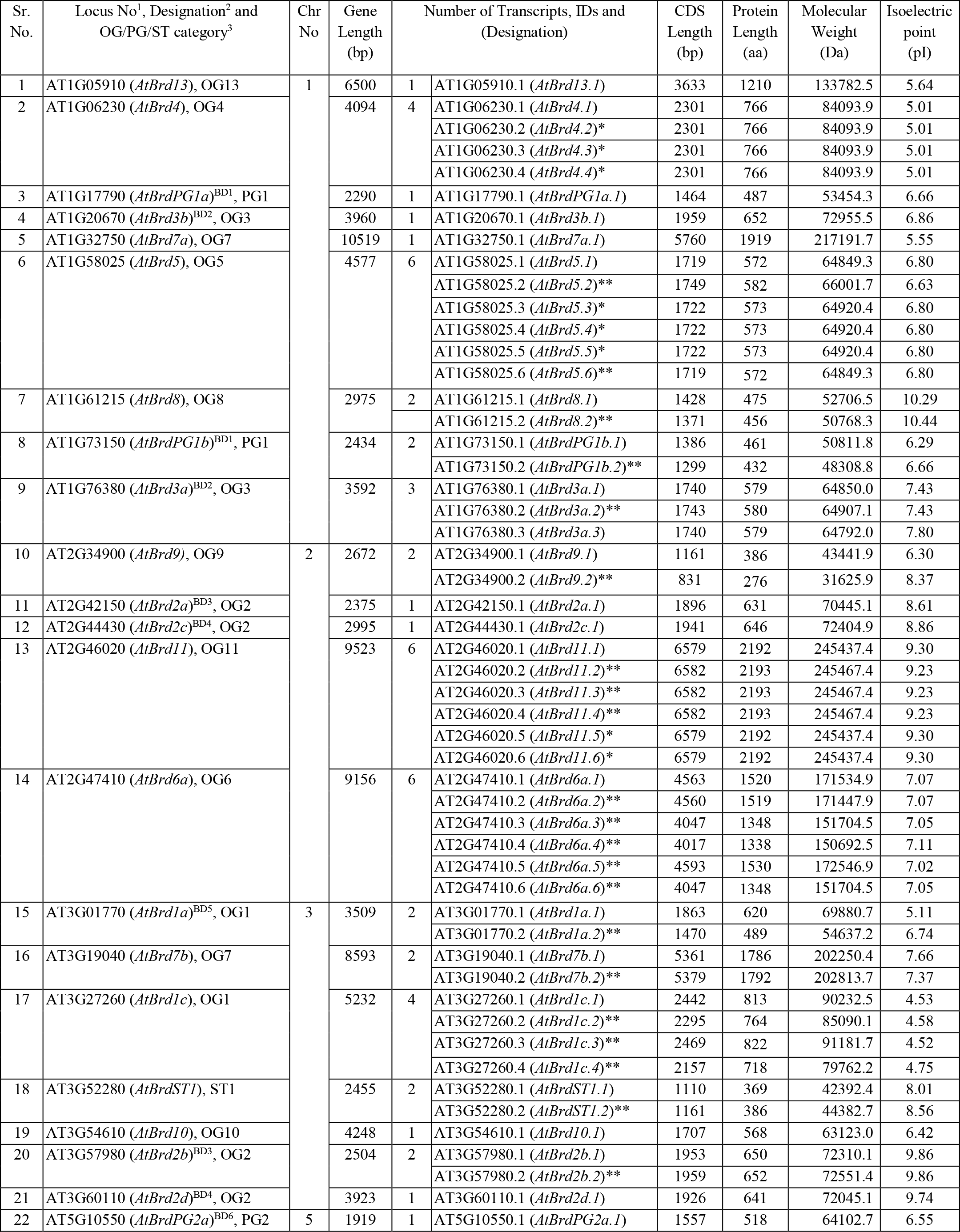
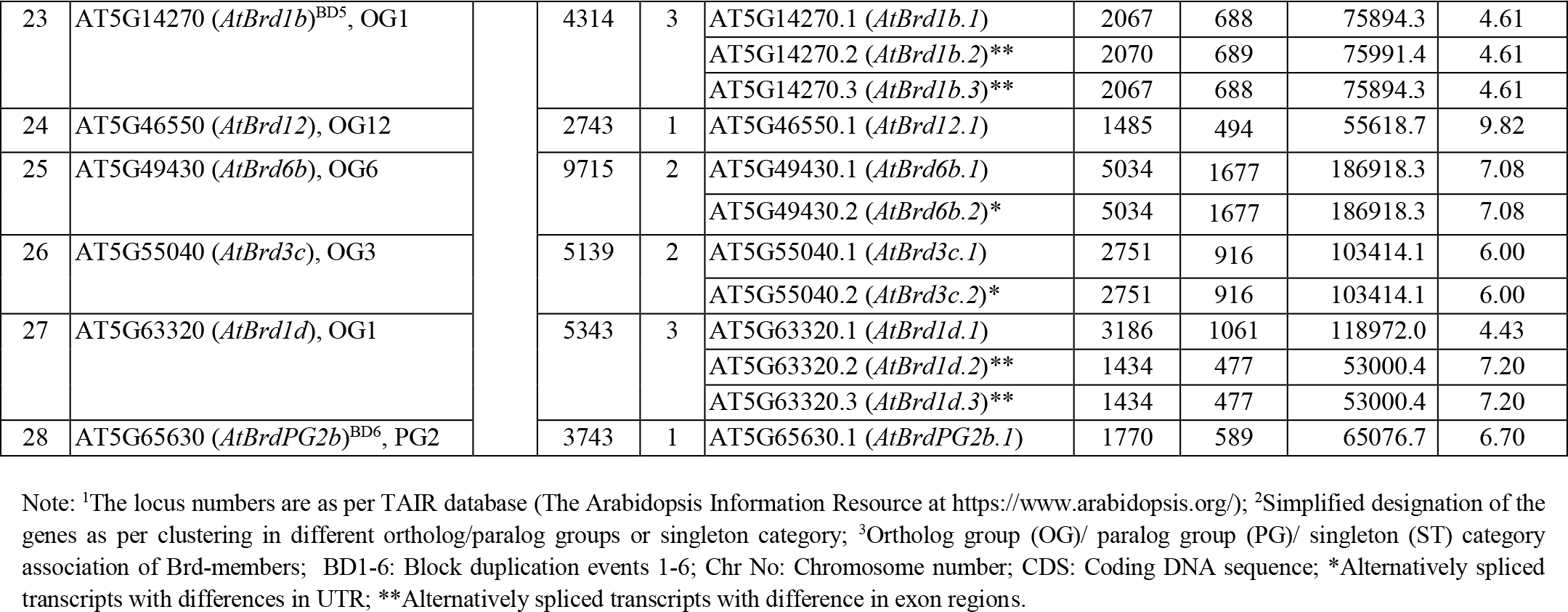
An overview of characteristics of 28 bromodomain-containing genes (*Brd*-genes) in *Arabidopsis thaliana* genome.

### 2.8 Multiple sequence alignment and phylogenetic analysis

Multiple sequence alignment of the bromodomain (BRD) region of AtBRDs and OsBRDs was done by ClustalX, and used for estimation of sequence divergence, and analysis of genetic relationships using neighbour-joining approach (Saitou and Nei, 1987) in Molecular Evolutionary Genetic Analysis X software (MEGAX, version 10.0.5, Kumar et al. 2018). Statistical analysis was carried out by bootstrap method (Felsenstein 1985). To identify the conserved residues in key elements of BRD-fold, the alignment was transformed into a sequence logo using TBtools (Chen et al., 2020). In a separate analysis, BRD regions of few human homologs containing single (UniProt accession numbers: Q9NR48, ASH1L; Q9NPI1, BRD7; Q9H0E9-2, BRD8B; P55201-1, BRPF1A; Q92830, GCN5L2; Q03164, MLL; Q13342, SP140; Q13263, TRIM28; Q9UPN9, TRIM33A; O15016, TRIM66; P51531, SMCA2; P51532, SMCA4) or two bromodomains (P25440, BRD2; Q15059, BRD3; O60885, BRD4; Q58F21, BRDT; Q6RI45, BRWD3; P21675, TAF1; Q9NS16, WDR9) were also included.

### 2.9 Homology modelling and comparison

The homology models of BRD-fold of several *A. thaliana* and *O. sativa* BRD proteins were generated at SWISS-MODEL workspace (http://swissmodel.expasy.org) using automated mode option and compared for structural differences. For identification of structural differences due to variations among BRD-folds, the homology models of BRD-folds of duplicate Brd-pairs or divergent Brd-members were superposed using structure comparison tools at SWISS-MODEL workspace.

### 2.10 Analysis of duplication events among plant genomes

The *Brd*-gene members from *A. thaliana* and *O. sativa* were analyzed for block and tandem duplication events at PLAZA (version 4.5) dicots and monocots databases (https://bioinformatics.psb.ugent.be/plaza/, Van Bel et al. 2018). InterPro identifier IPR001487 (bromodomain) was used to identify the chromosomal location of all *Brd*-genes (including duplicate-pairs), and represented using the Circle Plot tool available at PLAZA server. Additional 79 plant genomes including seven lower photosynthetic organisms, 27 monocots and 45 dicots (available at PLAZA monocots and dicots databases) were analyzed for prevalence of different duplication events (block, tandem, combined tandem + block events) leading to multiple *Brd*-genes.

## 3 Results

### 3.1 Diversity of Brd-members in *A. thaliana* and *O. sativa*: block and tandem duplications

Database analysis identified a total of 28 *Brd*-gene family members in *A. thaliana* and 22 members in *O. sativa* (Table 1 and 2). The *Brd*-genes displayed unequal chromosomal distribution in both *A. thaliana* (Chr1: 09, Chr3/Chr5: 07 each, Chr2: 05, Chr4: nil; Table 1, Figure 1A) and *O. sativa* (Chr2: 04, Chr3/Chr6/Chr8: 03 each, Chr1/Chr4/Chr7/Chr9: 02 each, Chr10: 01, Chr5/Chr11/Chr12: nil; Table 2, Figure 1B). The *Brd*-genes and encoded proteins in both the species showed extensive diversity in characteristics viz. number of alternative isoforms, molecular weight, isoelectric point (Table 1 and 2).

**Table 2.** An overview of characteristics of 22 bromodomain-containing genes (*Brd*-genes) in *Oryza sativa* genome.

| Sr. No. | Locus No<sup>1</sup>, Designation<sup>2</sup> and OG/PG/ST category<sup>3</sup> | Chr No | Gene Length (bp) | Number of Transcripts, Transcript IDs and (Designation) |  | CDS Length (bp) | Protein Length (aa) | Molecular Weight (Da) | Isoelectric Point (pI) |
| --- | --- | --- | --- | --- | --- | --- | --- | --- | --- |
| 1 | LOC_Os01g11580 (*OsBrd4a*)<sup>BD1</sup>, OG4 | 1 | 5988 | 2 | LOC_Os01g11580.1 (*OsBrd4a.1*) | 1068 | 355 | 39749.8 | 4.96 |
|  |  |  |  |  | LOC_Os01g11580.2 (*OsBrd4a.2*)** | 663 | 220 | 24655.7 | 4.42 |
| 2 | LOC_Os01g46040 (*OsBrdST1*)<sup>BD1</sup>, ST |  | 4046 | 1 | LOC_Os01g46040.1 (*OsBrdST1.1*) | 717 | 238 | 26206.1 | 6.95 |
| 3 | LOC_Os02g02290 (*OsBrd11*), OG11 | 2 | 10570 | 1 | LOC_Os02g02290.1 (*OsBrd11.1*) | 6603 | 2200 | 246212.0 | 9.07 |
| 4 | LOC_Os02g09920 (*OsBrdST2*), ST |  | 5584 | 1 | LOC_Os02g09920.1 (*OsBrdST2.1*) | 2940 | 979 | 110342.0 | 4.79 |
| 5 | LOC_Os02g15220 (*OsBrd4b*), OG4 |  | 7999 | 3 | LOC_Os02g15220.1 (*OsBrd4b.1*) | 1971 | 656 | 71476.3 | 10.00 |
|  |  |  |  |  | LOC_Os02g15220.2 (*OsBrd4b.2*)* | 1971 | 656 | 71476.3 | 10.00 |
|  |  |  |  |  | LOC_Os02g15220.4 (*OsBrd4b.3*)** | 1875 | 624 | 68946.6 | 10.20 |
| 6 | LOC_Os02g38980 (*OsBrd1*), OG1 |  | 5209 | 5 | LOC_Os02g38980.1 (*OsBrd1.1*) | 2145 | 714 | 78630.4 | 4.84 |
|  |  |  |  |  | LOC_Os02g38980.3 (*OsBrd1.3*)** | 1728 | 575 | 63133.5 | 5.62 |
|  |  |  |  |  | LOC_Os02g38980.4 (*OsBrd1.4*)** | 1701 | 566 | 62093.3 | 5.61 |
|  |  |  |  |  | LOC_Os02g38980.5 (*OsBrd1.5*)** | 1443 | 480 | 52786.3 | 6.67 |
|  |  |  |  |  | LOC_Os02g38980.6 (*OsBrd1.6*)** | 1302 | 433 | 48256.5 | 8.01 |
| 7 | LOC_Os03g03870 (*OsBrd3a*), OG3 | 3 | 6059 | 1 | LOC_Os03g03870.1 (*OsBrd3a.1*) | 1452 | 483 | 52387.9 | 8.43 |
| 8 | LOC_Os03g19340 (*OsBrd6*), OG6 |  | 16548 | 1 | LOC_Os03g19340.1 (*OsBrd6.1*) | 4881 | 1626 | 183000 | 6.64 |
| 9 | LOC_Os03g21450 (*OsBrdST3*), ST |  | 2605 | 1 | LOC_Os03g21450.1 (*OsBrdST3.1*) | 1677 | 558 | 59620.1 | 10.63 |
| 10 | LOC_Os04g53130 (*OsBrdPG3a*)<sup>TD1</sup>, PG3 | 4 | 3427 | 1 | LOC_Os04g53130.1 (*OsBrdPG3a.1*) | 1068 | 355 | 39412.4 | 4.81 |
| 11 | LOC_Os04g53170 (*OsBrdPG3b*)<sup>TD1-BD2</sup>, PG3 |  | 2150 | 1 | LOC_Os04g53170.1 (*OsBrdPG3b.1*) | 1371 | 456 | 50290.7 | 6.94 |
| 12 | LOC_Os06g04640 (*OsBrd9*), OG9 | 6 | 5300 | 3 | LOC_Os06g04640.1 (*OsBrd9.1*) | 1083 | 360 | 40658.1 | 7.05 |
|  |  |  |  |  | LOC_Os06g04640.2 (*OsBrd9.2*)** | 819 | 272 | 31690.2 | 6.19 |
|  |  |  |  |  | LOC_Os06g04640.3 (*OsBrd9.3*)** | 684 | 227 | 26418.0 | 6.32 |
| 13 | LOC_Os06g24870 (*OsBrd5a*)<sup>BD3</sup>, OG5 |  | 5765 | 3 | LOC_Os06g24870.1 (*OsBrd5a.1*) | 1137 | 378 | 40896.8 | 8.15 |
|  |  |  |  |  | LOC_Os06g24870.2 (*OsBrd5a.2*)* | 1137 | 378 | 40896.8 | 8.15 |
|  |  |  |  |  | LOC_Os06g24870.3 (*OsBrd5a.3*)* | 1137 | 378 | 40896.8 | 8.15 |
| 14 | LOC_Os06g43790 (*OsBrd7*), OG7 |  | 14210 | 1 | LOC_Os06g43790.1 (*OsBrd7.1*) | 5475 | 1824 | 206046 | 5.46 |
| 15 | LOC_Os07g32420 (*OsBrd12*), OG12 | 7 | 8036 | 3 | LOC_Os07g32420.1 (*OsBrd12.1*)* | 1455 | 484 | 53992.3 | 8.33 |
|  |  |  |  |  | LOC_Os07g32420.2 (*OsBrd12.2*)* | 1455 | 484 | 53992.3 | 8.33 |
|  |  |  |  |  | LOC_Os07g32420.1 (*OsBrd12.3*)* | 1455 | 484 | 53992.3 | 8.33 |
| 16 | LOC_Os07g37800 (*OsBrd8*), OG8 |  | 4117 | 3 | LOC_Os07g37800.1 (*OsBrd8.1*) | 1485 | 494 | 53351.1 | 10.54 |
|  |  |  |  |  | LOC_Os07g37800.2 (*OsBrd8.2*)** | 1407 | 464 | 50177.5 | 10.50 |
|  |  |  |  |  | LOC_Os07g37800.3 (*OsBrd8.3*)** | 1035 | 344 | 37209.5 | 8.15 |
| 17 | LOC_Os08g01794 (*OsBrd5b*)<sup>BD3</sup>, OG5 | 8 | 7581 | 2 | LOC_Os08g01794.1 (*OsBrd5b.1*) | 1773 | 590 | 65099.3 | 5.49 |
|  |  |  |  |  | LOC_Os08g01794.2 (*OsBrd5b.2*)* | 1773 | 590 | 65099.3 | 5.49 |
| 18 | LOC_Os08g09340 (*OsBrdPG3c*)<sup>BD2</sup>, PG3 |  | 2119 | 2 | LOC_Os08g09340.1 (*OsBrdPG3c.1*) | 1446 | 481 | 53638.1 | 6.63 |
|  |  |  |  |  | LOC_Os08g09340.2 (*OsBrdPG3c.2*)** | 1260 | 419 | 47311.1 | 9.65 |
| 19 | LOC_Os08g39980 (*OsBrd2*)<sup>BD4</sup>, OG2 |  | 3367 | 1 | LOC_Os08g39980.1 (*OsBrd2.1*) | 1983 | 660 | 68668.1 | 9.38 |
| 20 | LOC_Os09g33980 (*OsBrd13*)<sup>BD4</sup>, OG13 | 9 | 7027 | 1 | LOC_Os09g33980.1 (*OsBrd13.1*) | 3597 | 1198 | 133903 | 6.49 |
| 21 | LOC_Os09g37760 (*OsBrd3b*) |  | 5783 | 1 | LOC_Os09g37760.1 (*OsBrd3b.1*) | 1245 | 414 | 45957.8 | 9.84 |
| 22 | LOC_Os10g28040 (*OsBrd10*) | 10 | 6439 | 1 | LOC_Os10g28040.1 (*OsBrd10.1*) | 1536 | 511 | 56685.3 | 6.79 |

Note: <sup>1</sup>The locus numbers are as per RGAP database (Rice Genome Annotation Project at http://rice.uga.edu/); <sup>2</sup>Simplified designation of the genes as per clustering in different ortholog/paralog groups or singleton category; <sup>3</sup>Ortholog group (OG)/ paralog group (PG)/ singleton (ST) category association of Brd-members; BD1-4: Block duplication events 1-4; TD1: Tandem duplication event; Chr No: Chromosome number; CDS: Coding DNA sequence; Alternatively spliced transcripts with differences in UTR; \*\*Alternatively spliced transcripts with difference in exon regions.

**Figure 1:**
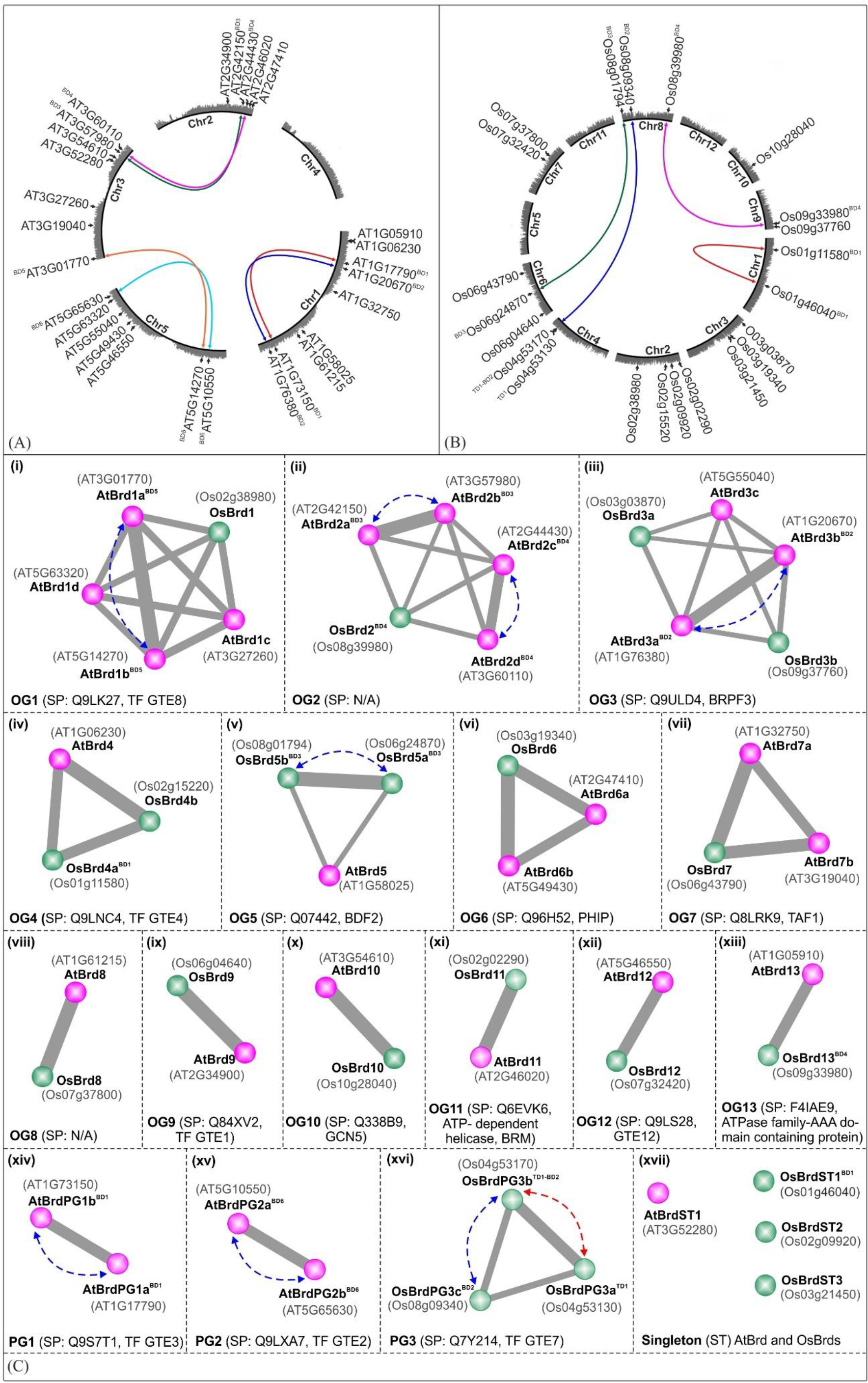
Circle-plot representation of chromosomal distribution of *Brd*-genes in *A. thaliana* (A) and *O. sativa* (B) genomes. The locus number designations are as per TAIR (for *AtBrds*) and RGAP (for *OsBrds*) databases. Colored connecting lines indicate the block-duplicated *Brd*-gene pairs, ‘Chr1-5/1-12’ indicate chromosome numbers, ‘BD’ and ‘TD’ indicate block and tandem duplication events. (C) Conserved ortholog groups (OGs), paralog groups (PGs), and singleton (STs) Brd-members in *A. thaliana* (purple circles) and *O. sativa* (green circles), as per analysis Orthology analysis at Orthovenn2 server (https://orthovenn2.bioinfotoolkits.net/). Blue and red colored dotted lines indicate the block-duplicated (BD) and tandem-duplicated (TD) Brds, while the functional information (based on SwissProt IDs) is indicated in the parenthesis (N/A indicates ‘No Hit’).

Syntenic analysis identified that in *A*. *thaliana* six *AtBrd*-gene pairs have originated due to block duplication events (BD1-BD6) (Figure 1A). Two intra-chromosomal block duplications were present in Chr1 (BD1: AT1G17790-AT1G73150 and BD2: AT1G20670-AT1G76380) and one in Chr5 (BD6: AT5G10550-AT5G65630). Remaining duplicated *AtBrd*-gene pairs involved inter-chromosomal block duplications viz. BD3 (Chr2-Chr3, AT2G42150-AT3G57980), BD4 (Chr2-Chr3, AT2G44430-AT3G60110) and BD5 (Chr3-Chr5, AT3G01770-AT5G14270) (Figure 1A). In *O. sativa* five *OsBrd*-gene pairs originated due to one tandem (TD1) and four block duplication events (BD1-BD4), of which one was an intra-chromosomal event in Chr1 (BD1: LOC_Os01g115800-LOC_Os01g46040) and three inter-chromosomal events (BD2: Chr4-Chr8, LOC_Os04g53170-LOC_Os08g09340; BD3: Chr6-Chr8, LOC_Os06g24870-LOC_Os08g01794 and BD4: Chr8-Chr9, LOC_Os08g39980-LOC_Os09g33980) (Figure 1B). In addition, LOC_Os04g53170 (of BD2) was also involved in a tandem duplication event leading to LOC_Os04g53130 in Chr4 (Figure 1B).

### 3.2 Orthologs and paralogs among *A. thaliana* and *O. sativa* Brd-members

Analysis of 28 AtBrds and 22 OsBrds at OrthoVenn2 server identified 13 ortholog groups (OG1-OG13), three paralog groups (PG1-PG3), and four singleton (ST) members (one AtBrd, three OsBrds) (Figure 1C). For subsequent description, the Brd-members were designated using a simplified scheme indicating their OG/PG association and duplication status, and based on five-components, 1) At/Os (species, At: *A. thaliana* and Os: *O. sativa*), 2) *Brd*/BRD (gene/ protein), 3) 1-13/ PG1-3/ ST1-3 (for OG/ PG/ STs), 4) a-d (multiple members in cluster), and 5) BD/TD (block/tandem duplication). For example, OG1 cluster contains one *OsBrd* (designated as *OsBrd1*) and four AtBrds (designated as *AtBrd1a*^BD5^, *AtBrd1b* ^BD5^, *AtBrd1c*, and *AtBrd1d*), of which two (*AtBrd1a*^BD5^and *AtBrd1b* ^BD5^) are outcome of block duplication, BD5 (Figure 1C-i). This scheme was useful to compare the evolutionary trend of Brd-members in the two species (Figure 1C).

The OGs showed variable representation of species-specific Brd-members and displayed different configurations relative to AtBrd-members viz. many-to-one (OG1, OG2, OG6, OG7; Figure 1C-i, 1C-ii, 1C-vi, 1C-vii), many-to-many (OG3, Figure 1C-iii), one-to-many (OG4, OG5, Figure 1C-iv, 1C-v), and one-to-one (OG8-OG13, Figure 1C-viii to 1C-xiii). Paralog groups PG1, PG2 were specific to *A. thaliana* (Figure 1C-xiv, 1C-xv), and PG3 was specific to *O. sativa* (Figure 1C-xvi). In certain OGs/PGs multiple Brd-members were due to species-specific block/tandem duplication events, viz. OG1 (BD5, Figure 1C-i), OG2 (BD3, BD4, Figure 1C-ii), OG3 (BD2, Figure 1C-iii), OG5 (BD3, Figure 1C-v), PG1 (BD1, Figure 1C-xiv), PG2 (BD6, Figure 1C-xv), PG3 (block tandem events in *O.* sativa, BD2 and TD1, Figure 1C-xvi). Interestingly, *OsBrd*-members of duplication events BD1 (*OsBrd4a-OsBrdST1*) and BD4 (*OsBrd2-OsBrd13*) clustered in different groups, suggesting relatively primitive events. The analysis identified conserved functions specific to different clusters viz. OG1 (transcription factor, TF GTE8), OG3 (bromodomain and PHD finger-containing protein 3, BRPF3), OG4 (transcription factor, TF GTE4), OG5 (bromodomain-containing factor, BDF2), OG6 (PH interacting protein, PHIP), OG7 (transcription initiation factor TF11D subunit 1, TAF1), OG9 (transcription factor, TF GTE1), OG10 (histone acetyltransferase, GCN5), OG11 (ATP dependent helicase, BRM a subunit of SWI/SNF multiprotein complex), OG12 (transcription factor, TF GTE 12), and OG13 (ATPase family-AAA domain containing protein). The paralog groups contained Brd-members with transcription factor functions viz. PG1 (TF GTE3, *A. thaliana*), PG2 (TF GTE2, *A. thaliana*) and PG3 (TF GTE7, *O. sativa*) (Figure 1C). The results also showed that duplication mediate *Brd-*gene copy number expansion was restricted to certain OG/PG groups, in both the species (*A. thaliana*: OG1-3 and PG1-2 and *O. sativa*: OG5, PG3) (Figure 1C).

### 3.3 Heterogeneity of *AtBrd* and *OsBrd*-members: impact of duplication and splicing events

The *AtBrd*-members showed considerable heterogeneity in length of the gene (1,919 - 10,519 bp), coding region (1,110 - 6,579 bp) and encoded proteins (369 - 2,192 AA) (Table 1), while the *OsBrds* displayed relatively higher variability (gene: 2,119 - 16,548 bp; coding region: 717 - 6,603 bp; protein: 238 - 2,200 AA) (Table 2), which was attributed to the length and number of exons, introns and 5′/ 3′-UTRs. The Brd-members in most OGs/PGs showed similarity in length and exon-intron organization in the two species. For example, OG2 members harbored 2-3 exons, while OG6 and OG7 contained extremely long *Brd-*genes with 17-24 exons (Figure 2).

**Figure 2:**
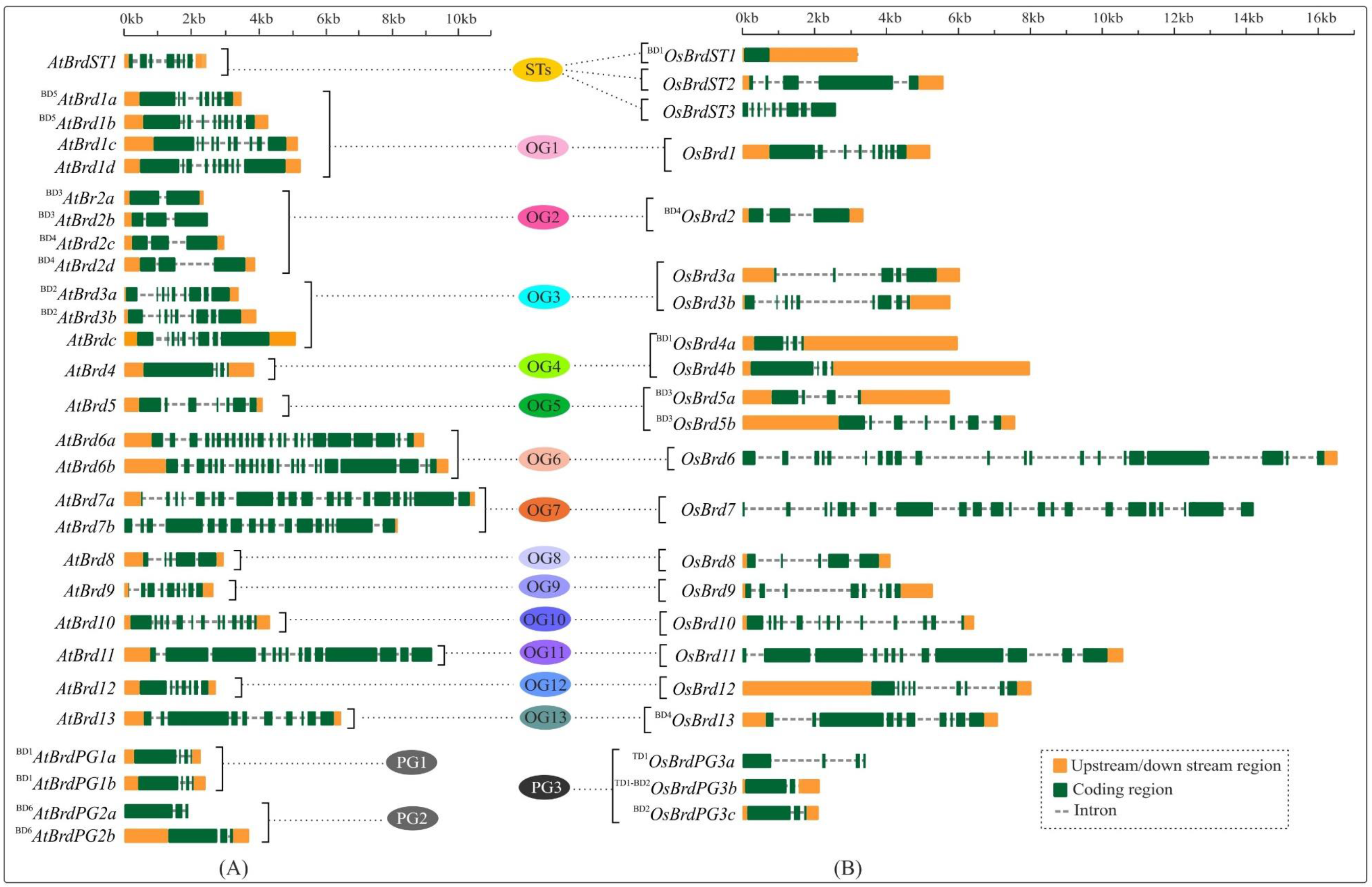
Comparison of gene structure and organization of *Brd*-genes from *A. thaliana* (A) and *O. sativa* (B), belonging to thirteen ortholog groups (OG1-13), three paralog groups (PG1-3), and singleton category (STs). The Brd-members specific to each group are arranged side-by-side for comparison. Different regions of genes are indicated by color codes (orange: upstream/downstream region including UTR, green: exons, and dashed line: intron). Scale on the top indicates the length in kilobase (kb), and designations ‘BD’ and ‘TD’ in the *Brd* names indicate the block or tandem duplication events.

Duplication events resulted in the diversity of the *Brd*-genes in both *A. thaliana* and *O. sativa*. The *AtBrd*-genes originated due to six block duplication events (BD1-BD6) displayed variations in length and organization of exons, introns and UTRs (Table 1, Figure 2A). Likewise, the five duplicated *OsBrd*-gene pairs due to one tandem and four block events displayed relatively higher heterogeneity than *AtBrds* (Table 2, Figure 2B). The *Brd*-members specific to certain OGs displayed species-specific duplication events viz. *AtBrd*-genes in OG1, OG2, and OG3 and *OsBrds* in OG5. Duplications resulted in higher heterogeneity in gene structure among the paralog *OsBrds* (PG3) than *AtBrds* (PG1, PG2) (Figure 2A and 2B).

In addition, AS-events also affected several *Brd*-genes in different OGs/PGs, which included around 60% *AtBrds* and 41% *OsBrd*-genes. In five ortholog groups (OG1, OG4, OG5, OG8, OG9), the *Brd*-genes of both the species showed AS, however the effects of events (on UTR/exon), and number of isoforms differed, with *OsBrd1* (OG1), *AtBrd5* (OG5) displaying highest number of transcripts (Supplementary Figure 1). In five OGs (OG2, OG3, OG6, OG7, OG11) AS-events were evident only among *AtBrd*-members including *AtBrd6a* (OG6) and *AtBrd11* (OG11) with maximum six isoforms, whereas in the OG12, only *OsBrd*-gene displayed AS-events (Supplementary Figure 1). One of the duplicated *Brd*-gene in PG1 (^BD1^*AtBrdPG1b, A. thaliana*), PG3 (^BD2^*OsBrdPG3c*, *O. sativa*) and *A. thaliana*-specific singleton *AtBrdST1*also accumulated variations to generate AS-transcripts. The AS-events affected UTRs in five genes (*AtBrd3c, AtBrd4*, *AtBrd6b*; ^BD3^*OsBrd5a*, ^BD3^*OsBrd5b*) coding regions in 13 genes (^BD5^*AtBrd1a*, *AtBrd1c*, ^BD3^*AtBrd2b*, *AtBrd7b*, *AtBrd8*, *AtBrd9*, ^BD1^*AtBrdPG1b*, *AtBrdST1*,; *OsBrd1*, ^BD1^*OsBrd4a*, *OsBRD8*, *OsBrd9*, ^BD2^*OsBrdPG3c*), and both in the remaining genes (Supplementary Figure 1). Interestingly, among the *Brd*-gene pairs affected by certain duplication events (*A. thaliana*: BD1, BD2, BD3 and *O. sativa*: BD1, BD2), only one of the copies displayed AS, whereas both the *Brd* copies generated by BD5 (*A. thaliana*) and BD3 (*O. sativa*) were affected by AS-events (Supplementary Figure 1). These results show that the OG-specific *Brd*-members and the duplicated gene-pairs seems to have evolved towards differential splicing patterns.

### 3.4 BRD-proteins showed heterogeneity in domain and motif organization

Apart from bromodomain (BRD), the AtBRD and OsBRD proteins harbored more than 25 other domains, including the most prominent extra-terminal (ET) domain (Figure 3). Certain domains were specific to AtBRDs (e.g. TLD, MDN1, Lys rich repeats, Figure 3A) and OsBRDs (e.g. PHD, WHIM1, Spo-VK, Med15, Asp/His rich repeats, Figure 3B). Broadly, AtBRD and OsBRD-members were divided into four types, a) containing only BRD, b) BRD + ET, c) BRD + other domains (other than ET), and d) BRD + ET + other domains. Notably, certain OGs with single Brd-members (OG9, OG10, OG12) showed conserved domain architecture, whereas, other OGs with single (OG8, OG11, OG13) and multiple Brd-members (OG1-OG7), including duplicated-Brds displayed domain differences (Figure 3). AtBRDs specific to *A. thaliana* PG1 and PG2 showed minor variations, whereas PG3-specific *(O. sativa*) ^BD2^OsBRDPG3C (duplicated by BD2 event) displayed a dual-BRD domain architecture (2^nd^ BRD overlapped with the ET domain) (Figure 3B). Furthermore, among the BRDs of the two species 15 conserved motifs (M1 - M15) were identified, of which M1 and M2 were most prevalent, and the duplicated members in the two species showed conserved signatures (Supplementary Table 1 and 2, Supplementary Figure 2).

**Figure 3:**
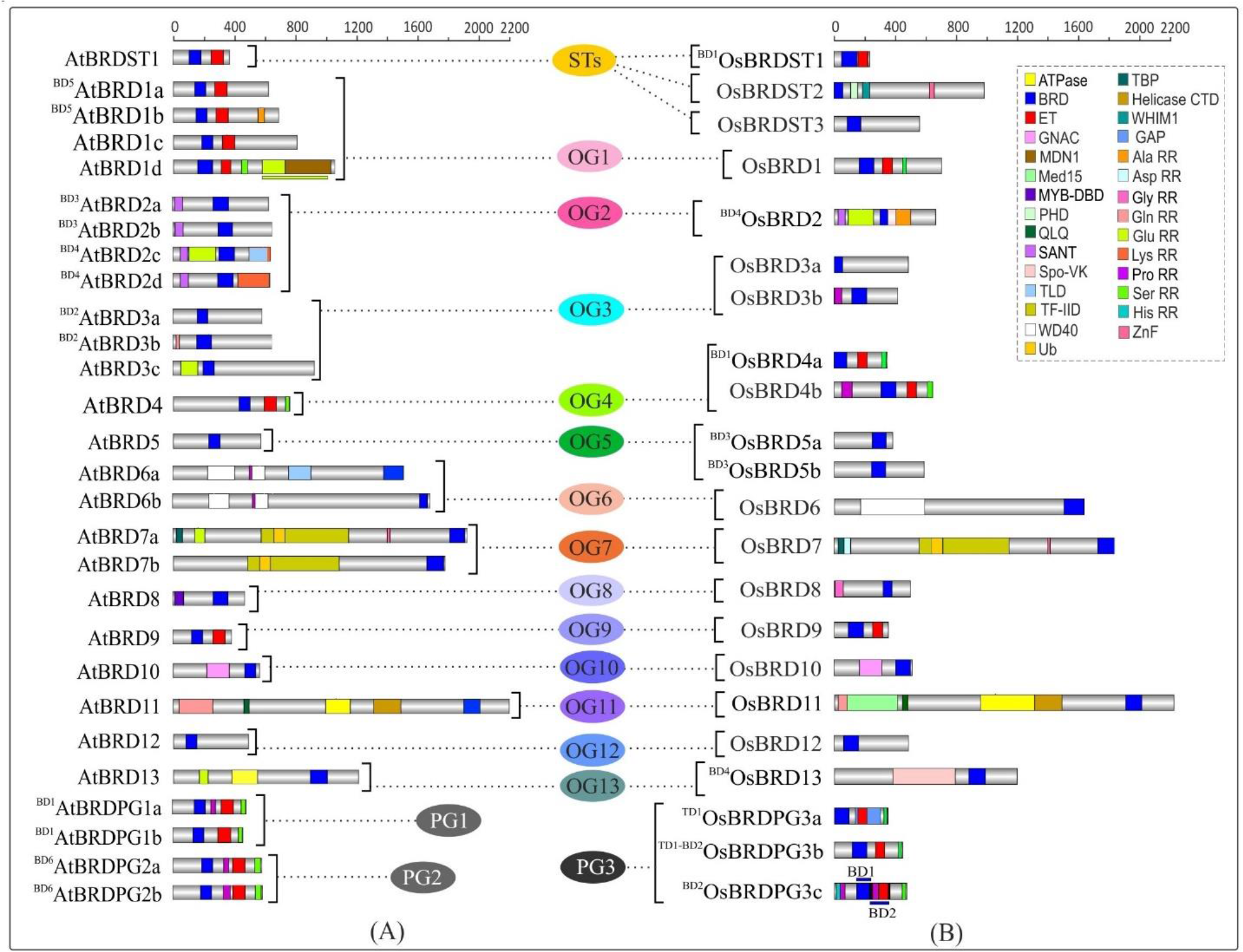
Comparison of domain architecture of BRD-proteins of *A. thaliana* (A) and *O. sativa* (B) belonging to thirteen ortholog groups (OG1-13), three paralog groups (PG1-3), and singleton category (STs). The Brd-members specific to each group are arranged side-by-side for comparison. Domains/important functional sites (as per CDD prediction) are shown by different color codes. In AtBRD27, OsBRD18 overlap of domains to adjacent regions is indicated by color-coded lines (above/below). Scale on the top indicates the protein length (amino acids) and designations ‘BD’ and ‘TD’ in the BRD names members indicate the block or tandem duplication events.

### 3.5 Duplications and AS-events affected domain architecture of BRDs

Duplication events affected the domain architecture of AtBRD and OsBRD-pairs. Four of the six block events (BD1, BD2, BD4, BD5) resulted in domain variations among the members of AtBRD-pairs than BD3 and BD6 events (Figure 3A). Among the OsBRD-duplicates, domain diversity was seen among members originated by tandem (TD1) and three block duplications (BD1, BD2, BD4), with substantial heterogeneity in BD2 and BD4 generated pairs (Figure 3B).

Coding-region (exon) specific AS-events also affected the domain diversity of several AtBRD and OsBRD-members (Figure 4). Eight AtBRDs from OG1, OG6, OG8-9, PG1 and ST1 displayed AS-mediated loss of certain domains (MYB-DBD: AtBRD8.2; MDN1: AtBRD1d.2, 1d.3; Ser RR: ^BD1^AtBRDPG1b.2) or N/C-terminal region (^BD5^AtBRD1a.2, AtBRD1c.2, 1c.4; AtBRD1d.2, 1d.3; AtBRD9.2; AtBRD6a.3, 6a.4, 6a.6; AtBRDST1.1; ^BD1^AtBRDPG1b.2) in alternative isoforms (Figure 4A). Likewise eight OsBRDs displayed AS-mediated loss of BRD (complete: ^BD1^OsBRD4a.2; partial: OsBRD8.2; OsBRD9.3), Ser RR (OsBRD1.6; OsBRD4b.3; OsBRDPG3c.2), and C-terminal truncation (OsBRD1.3, 1.4, 1.5, 1.6; OsBRD8.2, 8.3; OsBRD12.3; OsBRDPG3c.2) (Figure 4B). The AS-mediated loss/truncation of BRD was specific to three OsBRDs, and not observed in among AtBRDs. Both duplications and AS-events enhanced the diversity of BRD-members in two species.

**Figure 4:**
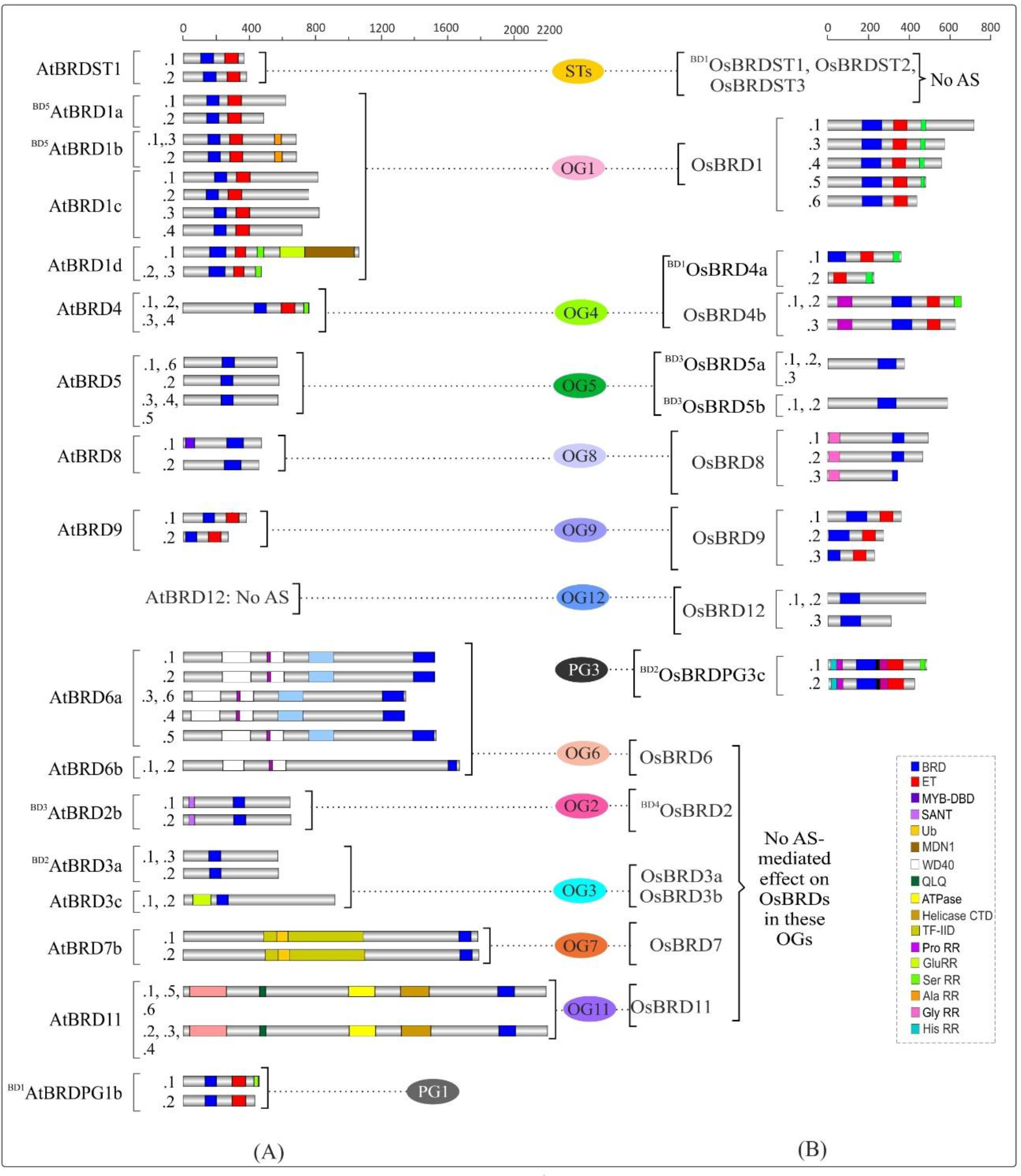
Alternative splicing (AS)-mediated changes in domain architecture of BRD isoforms of *A. thaliana* (A) and *O. sativa* (B) belonging to certain ortholog groups (OGs), paralog groups (PGs), and singleton category (STs). The Brd-members specific to each group are arranged side-by-side for comparison. Domains/important functional sites among different isoforms (constitutive: .1 and alternative: 0.2 to 0.6) are shown by different color codes. Scale on the top indicates the length (amino acids), and designations ‘BD’ and ‘TD’ in the BRD names indicate the block or tandem duplication events.

### 3.6 *Cis*-elements indicates responsiveness of *Brd*-genes to diverse intrinsic and extrinsic factors

The upstream regions of *Brd*-genes in both species contained *cis-*elements associated with diverse functions (Supplementary Figure 3 and 4), including response to light, stress conditions (abiotic: low temperature, anaerobic condition; biotic: wound, defense, elicitor-mediated activation), phytohormones (abscisic acid, auxin, salicylic acid, jasmonic acid, ethylene, gibberellin), and some physiological functions (Figure 5). While certain *Brd*-genes contained higher number of motifs for biotic stress (^BD4^*AtBrd2c*, ^BD4^*AtBrd2d*, *AtBrd3c*; ^BD4^*OsBrd2*, *OsBrd3b*, ^BD1^*OsBrd4a*, *OsBrd8*) and physiological functions (*AtBrdST1*, ^BD5^*AtBrd1b*, *AtBrd1c-1d*, ^BD3^*AtBrd2b*, ^BD2^*AtBrd3b*, *AtBrd12*, ^BD1^*AtBrdPG1a*, ^BD6^*AtBrdPG2a*; *OsBrd4b*, ^BD3^*OsBrd5a*, *OsBrd7*, *OSBrd10*, ^BD2^*OsBrdPG3c*), few lacked response motifs for phytohormone (*OsBrd3a*), abiotic stress (^BD1^*OsBrdST1*, *OsBrdST2*, ^BD4^*OsBrd2*) and light (*OsBrdST3*). Some *Brd*-genes displayed specific *cis-*elements viz. NON-box (*OsBrd4b*), motif1 (*OsBrd7*), and TATC-box (*OsBrd12*, ^BD3^*OsBrd5b*), MBSI (^TD1^*OsBrdPG3a* in *O. sativa*) (Supplementary Figure 3 and 4). Diversity of *cis-*elements indicate responsiveness of *Brd*-members towards diverse stimuli, with no conservation in different OGs/PGs (Figure 5A and 5B).

**Figure 5:**
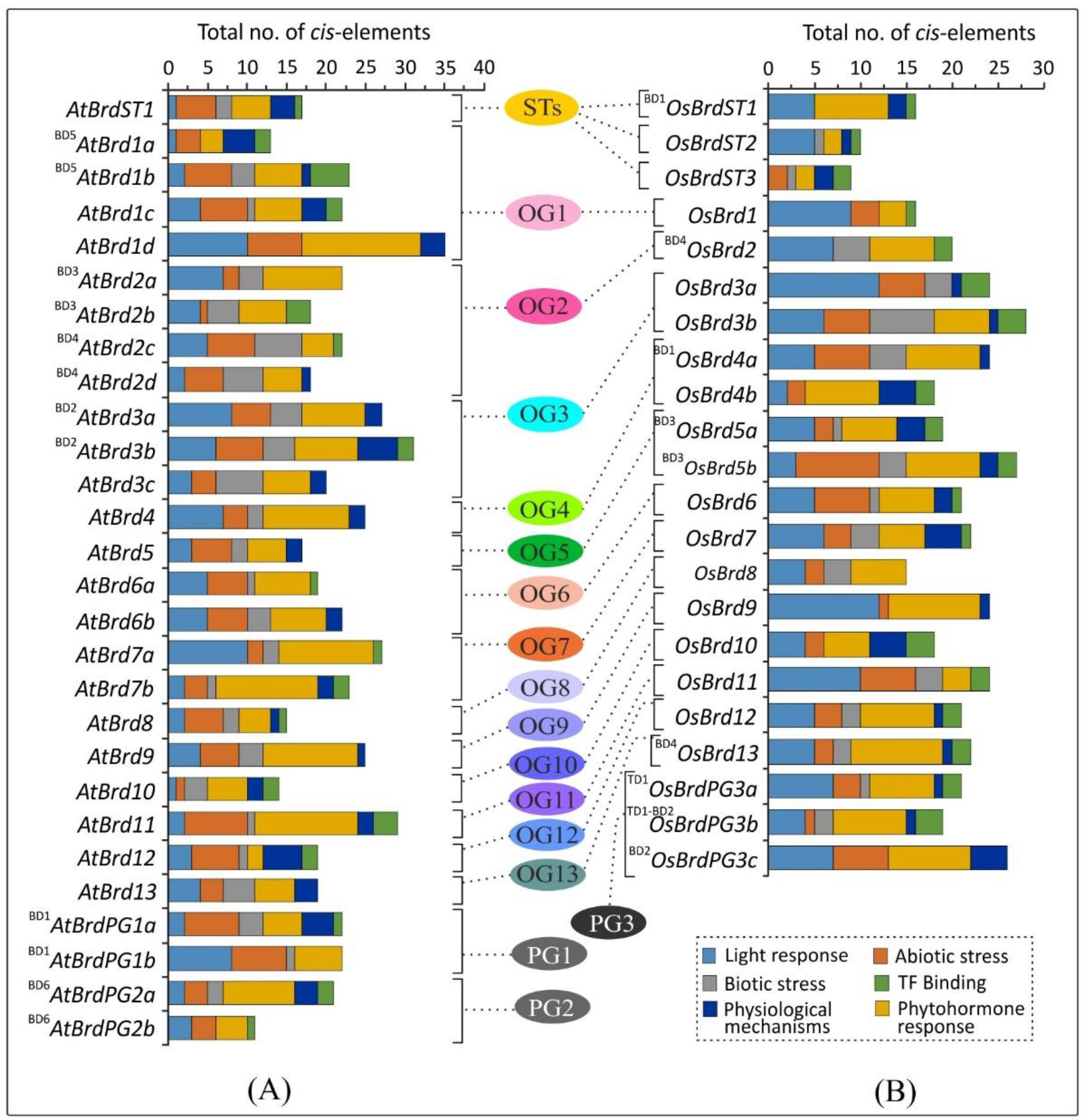
Diversity of *cis*-regulatory elements in the upstream region (−2000 bp) of *A. thaliana* (A) and *O. sativa* (B) *Brd*-genes, belonging to thirteen ortholog groups (OG1-13), three paralog groups (PG1-3), and singleton category (STs), as per analysis at PlantCARE database. The Brd-members specific to each group are arranged side-by-side for comparison, and different categories of *cis*-motifs are shown by different color code, and designations ‘BD’ and ‘TD’ in the names indicate block or tandem duplication events.

### 3.7 Duplication events affected the *cis*-element diversity and promoter structure

The tandem and block duplications affected the *cis-*element diversity among duplicated *Brd*-gene pairs in both the species. For example, *Brd*-pair ^BD5^*AtBrd1a-*^BD5^*AtBrd1b* (OG1) differed in *cis*-elements for light, abiotic and biotic stress (wound, defense), phytohormones (gibberellin, jasmonic acid) and physiological functions (meristem and endosperm-specific expression, circadian control), while ^BD1^*AtBrdPG1a*-^BD1^*AtBrdPG1b* (PG1) differed in elements for light, defense, abiotic stress, phytohormones (ethylene, gibberellin), TF-binding and physiological functions (Figure 5A, Supplementary Figure 3). *O. sativa* ^BD1^*OsBrd4a*-^BD1^*OsBrdST1* pair (OG4, ST) differed in *cis*-element copy number, and ^BD1^*OsBrdST1* also lacked elements for abiotic and biotic stress. OG5-specific ^BD3^*OsBrd5a*-^BD3^*OsBrd5b* displayed differences in elements for light, abiotic and biotic stresses, and certain physiological mechanisms (Figure 5B, Supplementary Figure 4). The duplications did not affected the length of promoter region of *AtBrd*-pairs, however substantial length differences were evident among most of the duplicate *OsBrd*-pairs (Figure 6). In addition, duplicated *AtBrd* and *OsBrd*-pairs displayed differences in the arrangement of TFBS (all duplicate pairs), repeat motifs (^BD2^*AtBrd3a*-^BD2^*AtBrd3b*; ^BD3^*AtBrd2a*-^BD3^*AtBrd2b*; ^BD1^*OsBrd4a*-^BD1^*OsBrdST1*, ^BD3^*OsBrd5a*-^BD3^*OsBrd5b*, ^BD4^*OsBrd2*-^BD4^*OsBrd13*), and CpG islands (^BD3^*AtBrd2a*-^BD3^*AtBrd2b*; ^BD5^*AtBrd1a*-^BD5^*AtBrd1b*; All *OsBrd*-pairs) (Figure 6). Overall, the duplication event seems to have affected the promoters, and responsiveness of the *Brd*-pairs towards diverse intrinsic/extrinsic factors.

**Figure 6:**
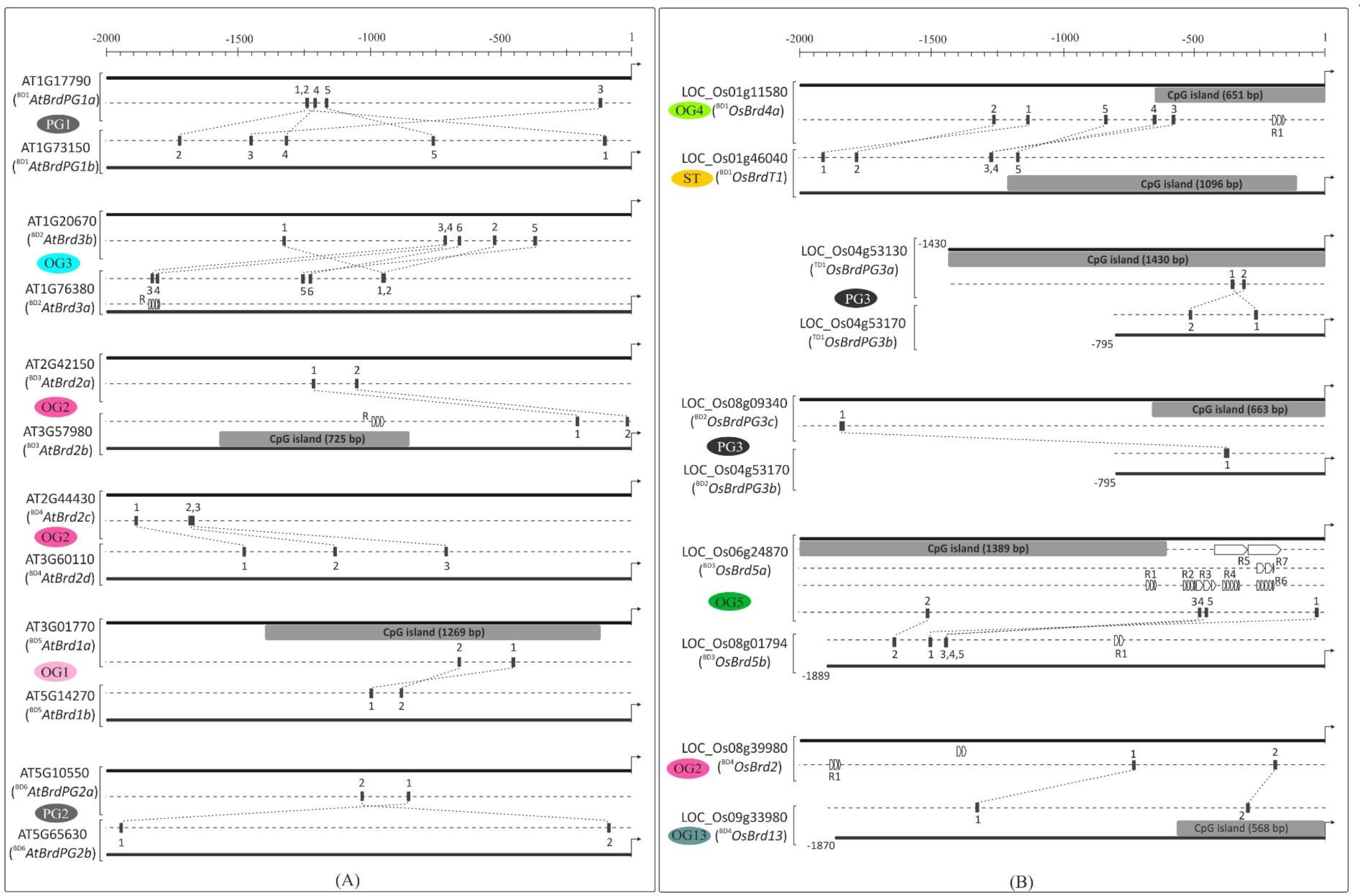
Analysis of upstream region (−2000 bp) of duplicated *Brd*-genes of *A. thaliana* (A) and *O. sativa* (B) on PlantPAN3.0 database, for difference in CpG islands (grey boxes), transcription factor binding sites (TFBS, indicated with numerals w.r.t different genes) and repetitive motifs (R). The designation (and locus number) of Brd-members, association with ortholog (OG)/paralog group (PG) or and singleton category (ST) is indicated. Scale on the top indicates the length of upstream region (bp), arrow towards right indicates translation start site, and designations ‘BD’ and ‘TD’ in the names indicate block or tandem duplication events.

### 3.8 *AtBrd* and *OsBrd*-genes showed tissue-and stress specific expression differences

RT-qPCR analysis of duplicate *Brd*-pairs (*AtBrd*: 4-pairs; *OsBrds*: 5 pairs) in seedlings tissue showed difference in basal transcript levels and response to salinity. Among the *AtBrd*-duplicates, ^BD1^*AtBrdPG1b*, ^BD3^*AtBrd2b*, and ^BD6^*AtBrdPG2b* showed higher transcript levels than corresponding duplicate members, while the BD4-pair showed comparable levels (Figure 7A, top panel). In response to salt stress, four *AtBrds* (^BD1^*AtBrdPG1a*; ^BD1^*AtBrdPG1b*; ^BD4^*AtBbrd2c*; ^BD4^*AtBrd2d*; ^BD6^*AtBrdPG2b*) were up-regulated (~2-6-fold), *AtBrd2a* was down-regulated and two (^BD6^*AtBrdPG2a*; ^BD3^*AtBrd2b*) remained unaffected (Figure 7A, bottom panel). In rice seedlings among the *OsBrd*-pairs, ^BD1^*OsBrd4a*, ^BD4^*OsBrd13*, ^BD3^*OsBrd5a*, ^BD2^*OsBrdPG3c* showed relatively higher basal transcript levels that the duplicate member (Figure 7B, top panel). Under salt stress, five *OsBrds* showed up-regulated (^BD1^*OsBrdST1*, ^BD3^*OsBrd5a*, ^BD3^*OsBrd5b*, ^TD1-BD2^*OsBrdPG3b*, ^BD2^*OsBrdPG3c*), ^BD4^*OsBrd2* was down-regulated, and three (^BD1^*OsBrd4a*, ^BD4^*OsBrd13*, ^TD1^*OsBrdPG3a*) showed no significant change (Figure 7B, bottom panel).

**Figure 7:**
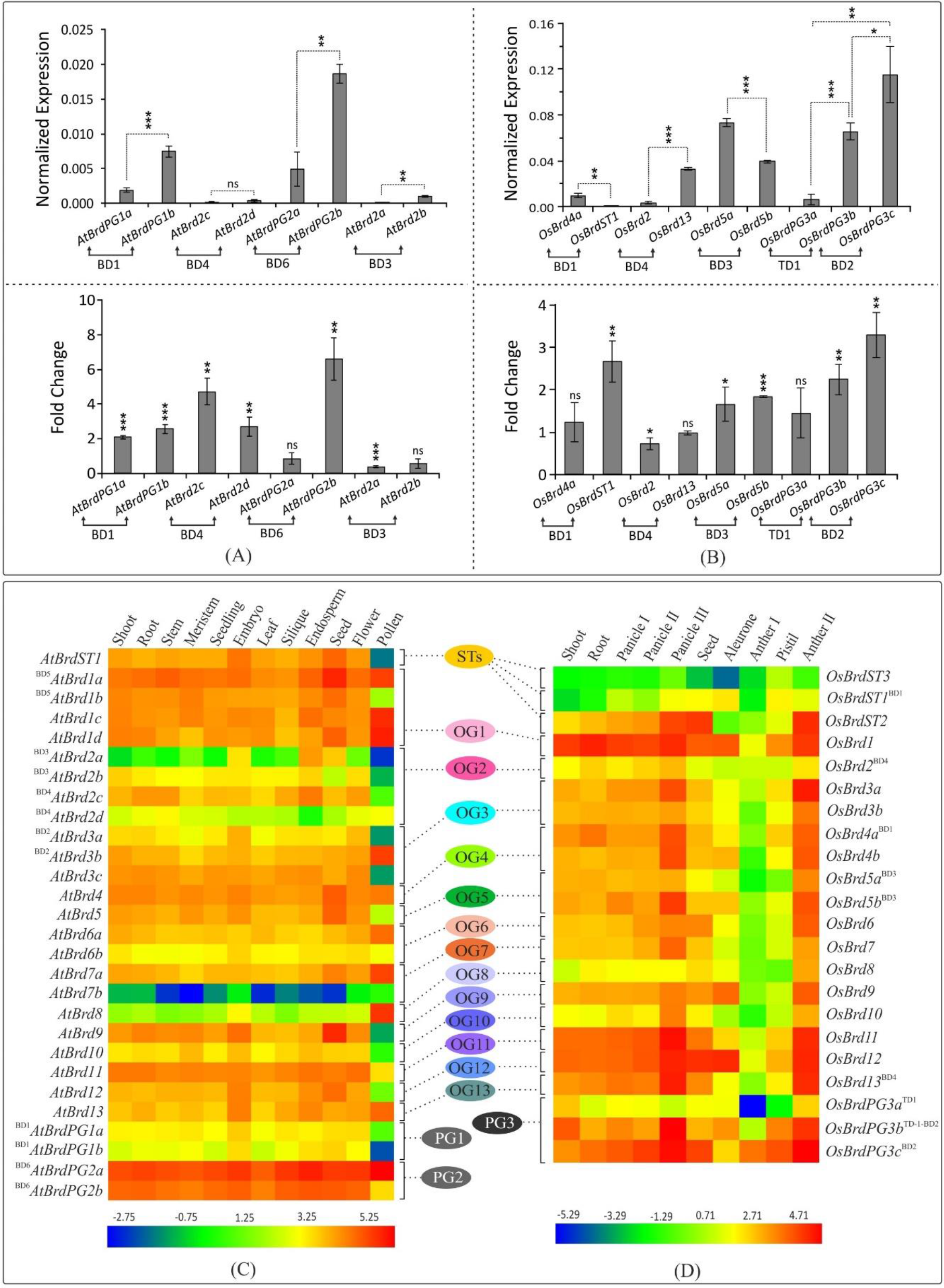
Transcript analysis of *Brd*-gene. RT-qPCR analysis of transcript levels four duplicated *AtBrd*-pairs (A) and five *OsBrd*-pairs (B) in seedling tissues (top panels), and in response to salt stress (NaCl, 150 mM, bottom panels), using reference genes (*AtActin*: AT3G10920; *OselF1α*: LOC_Os03g08010). Designations ‘BD’ and ‘TD’ in the names indicate block or tandem duplication event. The analysis was carried out in triplicate, data is represented as mean ±SD, and statistical significance is indicated by *(*p* <0.05), **(*p* <0.01), ***(*p* <0.001). Heatmap-based analysis of RNA-Seq data for tissue-specific expression pattern of *Brd*-genes of *A. thaliana* C) and *O. sativa* (D), belonging to thirteen ortholog groups (OG1-13), three paralog groups (PG1-3) and singleton category (STs). Details of tissues are indicated on the top, *Brd*-genes are indicated on the sides with designations ‘BD’ and ‘TD’ indicating block or tandem duplication event, and a color gradient scale indicates the expression level (blue: low levels; red: high)..

Further analysis of RNA-Seq data showed tissue-specific abundance patterns of *AtBrd*-genes. In general, *AtBrd*s from OG8, OG7 (*AtBrd7b*), OG2 (two genes: ^BD3^*AtBrd2a*, ^BD4^*AtBrd2d*) and PG1 showed lower transcript levels, while those from PG2, OG1, OG3-4, OG9 and PG3. Genes ^BD5^*AtBrd1a* and ^BD6^*AtBrdPG2a* showed high transcript levels in most tissues while *AtBrd7b* showed the lowest (Figure 7C). Pollen tissue displayed abundance of seven *AtBrds* (^BD5^*AtBrd1a*, *AtBrd1c*, *AtBrd1d*, ^BD3^*AtBrd3b*, *AtBrd7a, AtBrd8*, ^BD6^*AtBrdPG2a*), while many others showed lowest levels. Substantial tissue-specific differences were observed among members of two duplicate pairs, ^BD3^*AtBrd2a*-^BD3^*AtBrd2b* and ^BD4^*AtBrd2c*-^BD4^*AtBrd2d* (Figure 7C). In *O. sativa*, *OsBrds* from clusters OG8, OG10 and PG3 (^TD1^*OsBrdPG3a*) and two STs (^BD1^*OsBrdST1, OsBrdST3*) showed low transcript levels, while members from OG1, OG11-12 and PG3 (^BD2^*OsBrdPG3c*) were abundant in most tissues. In general, *OsBrds* showed low levels in anther I tissue and highest in panicle II and anther II stages (Figure 7D). Most *OsBrd*-duplicates displayed tissue-specific differences, with maximum variation in ^BD1^*OsBrd4a*-^BD1^*OsBrdST1* and ^BD4^*OsBrd2*-^BD4^*OsBrd13* pairs (Figure 7D). Together these result show that the *Brd*-duplicates have evolved for differential response towards intrinsic/extrinsic factors.

### 3.9 Sequence divergence and key conserved sites in bromodomain (BRD) region of *Brd*-genes

The bromodomain (BRD) region showed more length variation among OsBRDs (range: 57-133 AA) than AtBRDs (range: 94-133 AA), particularly due to two OsBRDs, OsBRD3a, OsBRDST2 that harbored long N-terminal deletion leading to exceptionally small BRD-regions (Supplementary Figure 5). Also, three-pairs of AtBRDs (^BD5^AtBRD1a-^BD5^AtBRD1b; ^BD3^AtBRD2a-^BD3^AtBRD2b; ^BD4^AtBRD2c-^BD4^AtBRD2d;) and OsBRDs (^BD1^OsBRD4a-^BD1^OsBRDST1, ^TD1-BD2^OsBRDPG3b-^BD2^OsBRDPG3c; ^BD4^OsBRD2-^BD4^OsBRD13) showed indel variations (Supplementary Figure 5). (Supplementary Figure 5). Several conserved residues, similar to human BRDs, were identified in the characteristic BRD-fold elements viz. αZ-helix (Leu-34, Ile/Leu-37, Leu-38, Leu/Ile-41), ZA-loop (Phe-52; Pro-55, −73; Val-56; Asp-65; Tyr-66; Ile-70; Met-74), αA-helix (highly conserved Asp-75; Leu-76, −83; Thr-78; Ile-79), small AB-loop (conserved Tyr-96), αB-helix (invariant Asp-105 and Phe-102, −110; Leu-108; Asn-112, −117; Tyr-116), and αC-helix (Val-123; Tyr-127; Met-129; Leu-133; Phe-137). Plant-specific signatures were also evident in ZA loop (Asp-57), αB-helix (Val-106, Thr-109, Ala-113, Met-114) and αC (Pro-118, Ala-130, Trp-141) (Figure 8A). Most of these key sites were also conserved among the BRD-duplicates, however AtBRD-pairs displayed variations from one (Ile →Val, in ZA-loop, OG1, PG1, PG2) to seven sites (^BD4^AtBRD2c-^BD4^AtBRD2d, OG2) (Figure 8B, top panel). With more heterogeneity, the OsBRD-pairs showed up to 20 variable sites (^BD4^OsBRD2-^BD4^OsBRD13) and loss of αZ-helix in ^BD1^OsBRD4a (^BD1^OsBRD4a-^BD1^OsBRDST1) (Figure 8B, bottom panel). Such variations can alter the interactions of the BRD-fold with chromatin.

**Figure 8:**
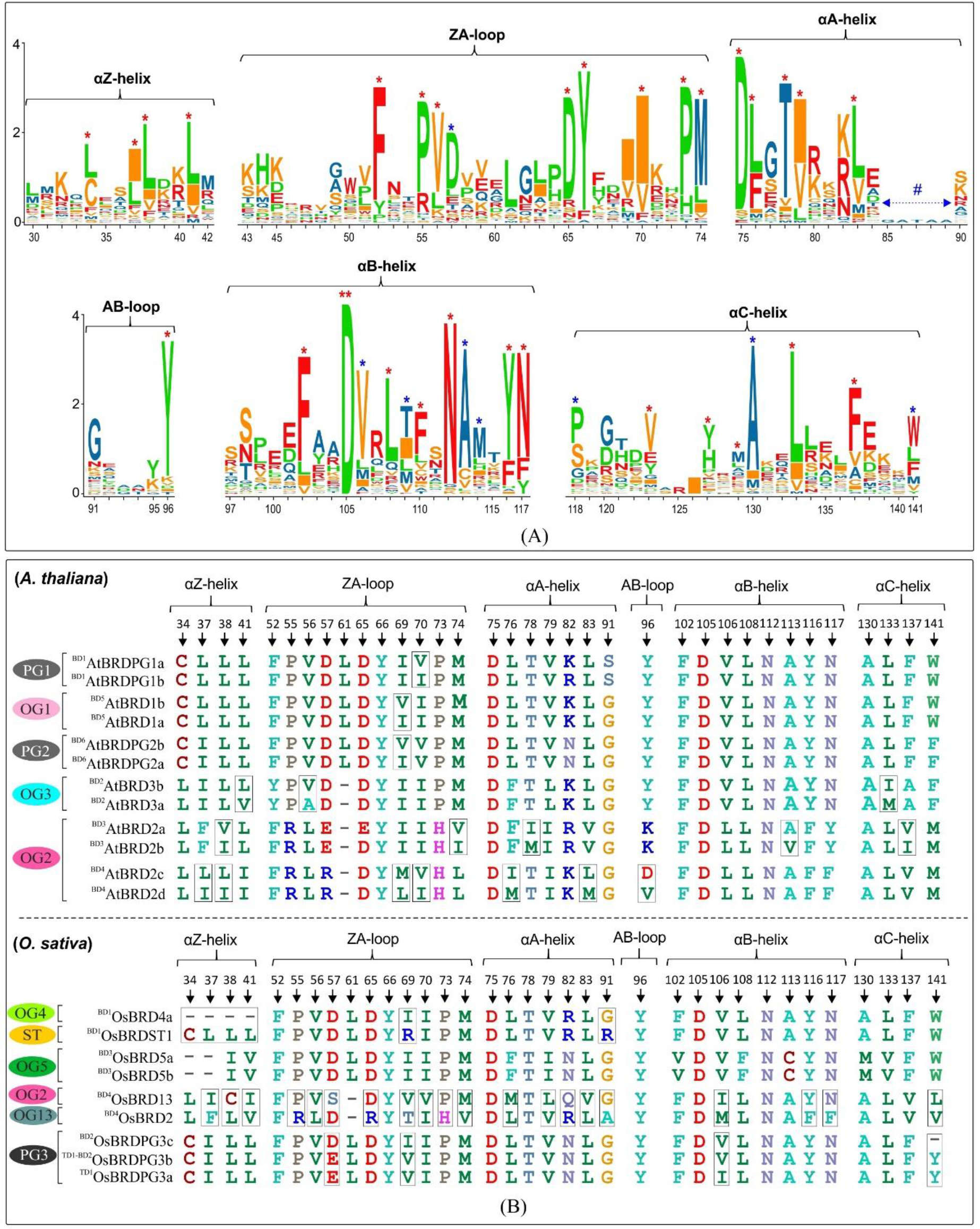
(A) Sequence logo analysis of the BRD-region of 28 AtBRD and 22 OsBRD-homologs, indicating conserved residues in the key BRD-fold elements (helices: αZ, αA, αB, αC; loops: ZA, AB, BC). ‘*’ indicates conserved sites (red ‘*’: conserved residues in human BRDs; blue ‘*’: conserved sites in At- and OsBRDs), and ‘**’ indicates an invariant residue. Site/positions of residues (as per the alignment in Supplementary Figure 5) are indicated the x-axis. (B) Comparison of conserved sites in key elements of BRD-fold among the duplicated BRDs of *A. thaliana* (top panel) and *O. sativa* (bottom panel). The association of duplicate-BRDs to different OGs/PGs or ST category is indicated, designations ‘BD’ and ‘TD’ indicate type of duplication event, and variations at key positions among the duplicates are indicated by rectangular boxes.

Cluster analysis based on BRD-region placed the 50 BRD sequences from two species into six clusters (I - VI) (Figure 9A). The site intra-group variability ranging from 21% to 67.3% (II and VI), while the intergroup variability ranged from 60.4% (I/IV) to 77.8% (III/VI). Different clusters/sub-clusters represented BRD-members specific to different OGs/PGs (Figure 2). Largest cluster I was divided into five sub-clusters: IA (OG4, PG1, PG2, PG3), IB (OG1, OG12), IC (^BD1^OsBRDST1), ID (OG9, AtBRDST1), IE (OG5). Other clusters also displayed similar trend viz. II (OG10), III (OG7, OG13), IV (OG3, OsBRDST3), V (OG2, OG8), and VI (OG6, OG11, OsBRDST2). The BRD-regions of all AtBRD-duplicate pairs (events: BD1 to BD6), and three OsBRD-pairs (events: TD1, BD2, BD3) grouped together in respective clusters indicative of less divergence (Figure 9A). On the contrary, members of two BD-pairs (^BD1^OsBRD4a-^BD1^OsBRDST1; ^BD4^OsBRD2-^BD4^OsBRD13) were clustered differently (IA, IC and III, V) indicating high divergence (Figure 9A). Consistency in the BRD-based clustering and the OG/PG grouping suggest similar divergence of the domain vis-à-vis total protein. Analysis with human single/dual BRD-regions identified clusters/sub-clusters specific to plants (GIA - IE, GII) and human sequences (GVII, IF – IH) (Figure 9B). Interestingly, the two domains of human dual BRD-members clustered with At/OsBRDs from different groups. For example, BRD(1) of BRD2-4, BRDT (sub-cluster IH) was close to IG (AtBRD5, ^BD3^OsBRD5a-^BD3^OsBRD5b), and BRD(2) sub-cluster (IE) was close to plant-specific sub-clusters (ID, IC, IB). Also, two BRD-domains of human WDR9 displayed high divergence and placed in different clusters (IG, GII) (Figure 9B).

**Figure 9:**
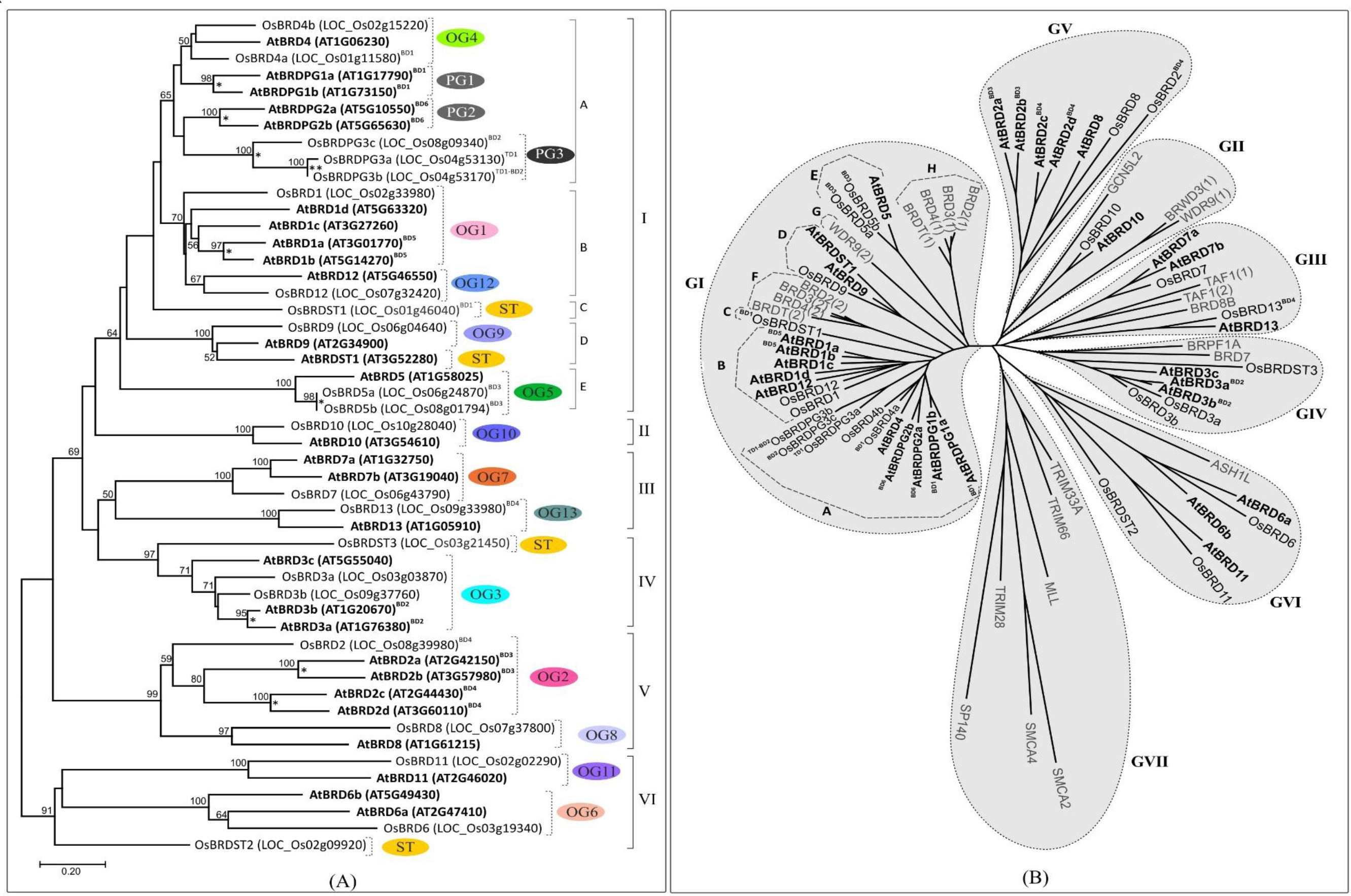
(A) Neighbor-joining phylogenetic tree based on the BRD-regions of Brd-homologs (including the block/tandem duplicates) of *A. thaliana* and *O. sativa*, generated by MEGA-X software. Major clusters ate indicated by Roman numerals (I-VI), while sub-clusters are shown by letters (A-E). The ortholog group (OG), paralog group (PGs) or singleton category (ST) is shown for different clusters/sub-clusters. Numbers at the nodes indicate bootstrap values (in %, for 500-replicates), and taxa names (AtBRDs: bold font, OsBRDs: regular font) include Brd-designation used, locus numbers (in parenthesis), and type of duplication events (BD: block duplication; TD: tandem duplication). (B) Radiation tree of the BRD-regions of *A*. *thaliana*, *O*. *sativa*, some representative human BRD-homologs (BRD2-4, BRD8B, BRDT, WDR9, TAF1, BRWD3, BRPF1A, BRD7, GCN5L2, ASH1L, TRIM33A, TRIM66, MLL, SMCA2, SMCA4, TRIM28, SP140). Seven major groups (GI-GVII) and subgroups (A-H) are indicated, where different BRDs are shown in different font style (AtBRDs: bold font; OsBRDs: regular font; human BRDs: grey font). Designation ‘BD’ and ‘TD’ indicated block and tandem duplication events, while numerals in parenthesis (1/2) indicate two domains of the dual-BRD-containing homologs.

### 3.11 Heterogeneity mediated structural variations in the bromodomain-fold (BRD-fold)

The BRD-fold is comprised of four α-helices (αZ, αA, αB, αC) and three loops (ZA, AB, BC) (Figure 10A), which were affected by both length/sequence variations in At- and OsBRDs (Supplementary Figure 5). Conserved BRD-fold was observed for several At/OsBRDs (Figure 10B), however sequence divergence affected prominent structural features viz. truncated αZ-helix due to N-ter deletion (^BD3^OsBRD5a, ^BD3^OsBRD5b, Figure 10C), an extended region before αZ-helix (AtBRD6a, OsBRD6), variation in ZA-loop (AtBRD6a) (Figure 10D), an extra α-helix after αC-helix due to long C-ter region (ATBRD7b, OsBRD7, Figure 10E), and complete loss of αZ-helix and ZA-loop (OsBRD3a, OsBRDST2, Figure 11F). Superposition of duplicated-BRD models revealed structural variations in the BRD-fold, including minor structural variations in AtBRD-pairs with low divergence (^BD3^AtBRD2a-^BD3^AtBRD2b, Figure 10G; ^BD5^AtBRD1a-^BD5^AtBRD1b, Figure 10H), loss of αZ-helix in ^BD1^OsBRD4a (^BD1^OsBRD4a-^BD1^OsBDRST1, Figure 10I), and variations in αZ, αC and BC loop (^BD4^OsBRD2-^BD4^OsBRD13, Figure 10J). Such structural variations might alter the characteristics and BRD-associated functions of duplicate-members.

**Figure 10:**
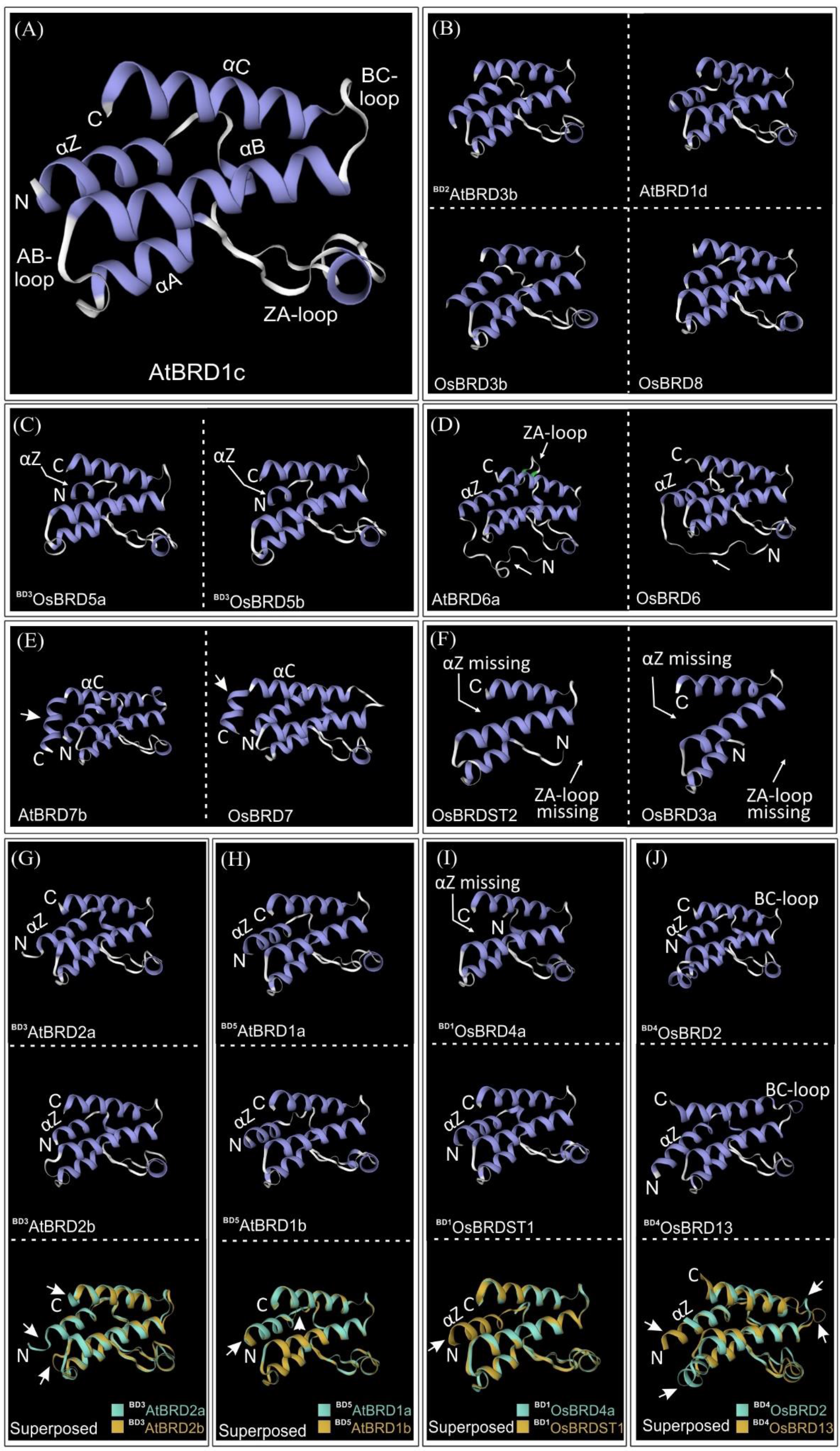
Homology models of BRDs-folds of Brd-homologs of *A. thaliana* and *O. sativa* generated at SWISS-MODEL workspace. (A) AtBRD1c with key BRD-fold elements (α-helices: αZ, αA, αB, αC; loops: ZA, AB and BC), (B) ^BD2^AtBRD3b, ArBRD1d, OsBRD3b and OsBRD8, C) BD3OsBRD5a and ^BD3^OsBRD5b, (D) AtBRD6a and OsBRD6, (E) AtBRD7b and OsBRD7, (F) OsBRDST2 and OsBRD3a. Structural superposition of BRD-models of duplicate BRD-pairs (shown in different colors: (G) ^BD2^AtBRD2a-^BD2^AtBRD2b, (H) ^BD5^AtBRD1a-^BD5^AtBRD1b, (I) ^BD1^OsBRD4a-^BD1^OsBRDST1, and (J) ^BD4^OsBRD2-^BD4^OsBRD13. Variations due to sequence/length heterogeneity or duplication events are indicated by arrows.

**Figure 11:**
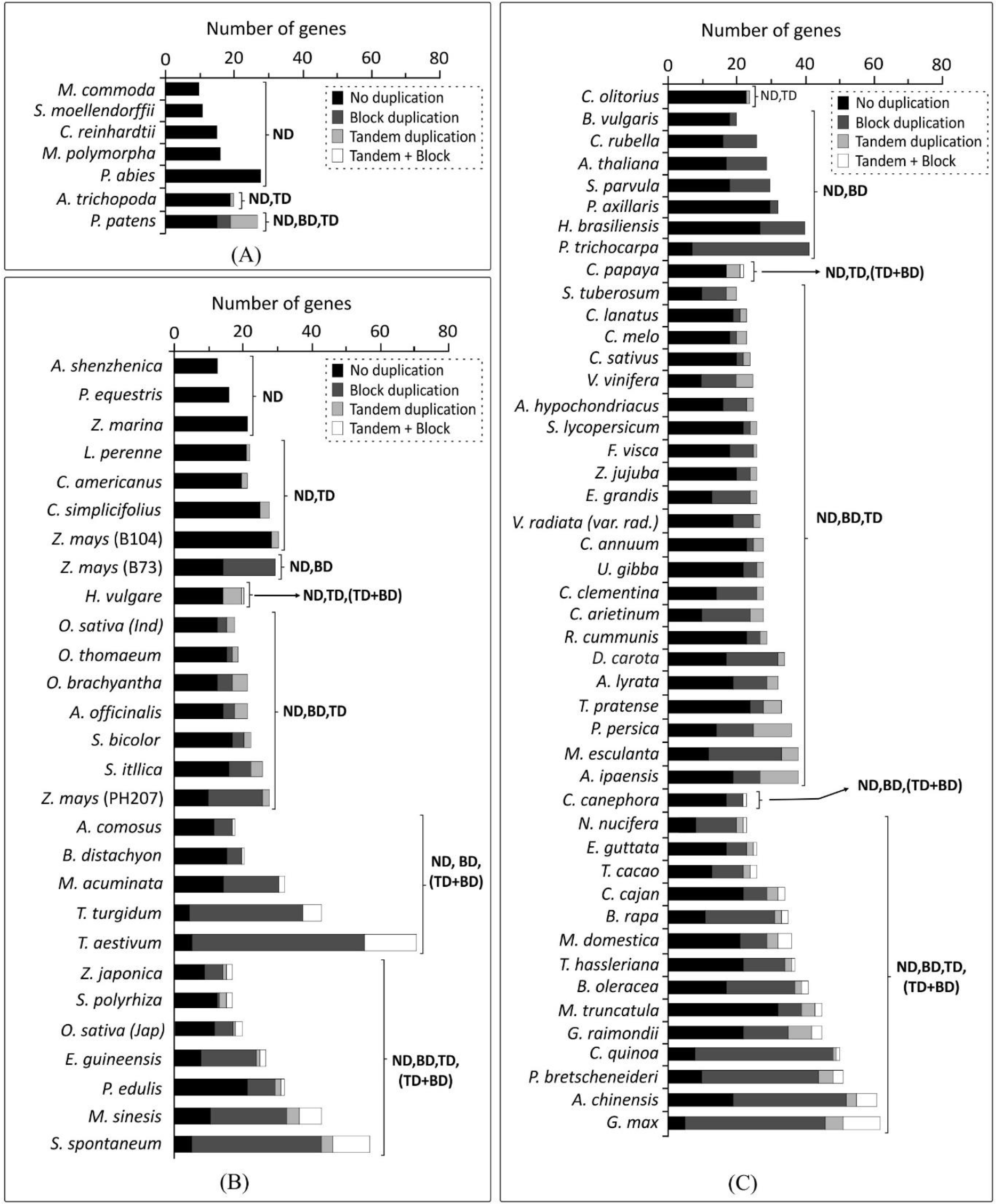
Comparative assessment of duplication events affecting *Brd*-gene copies among plant genomes: (A) lower photosynthetic organisms, (B) monocots, and (C), as per analysis at PLAZA database (version 4.5). Types of duplication events are indicated by different grey shades and designations ND (no duplication), TD (tandem duplication), BD (block duplication), and TD + BD (combine tandem and block duplication).

### 3.12 Duplication events affected the *Brd*-gene numbers among higher plants

Based on the results obtained in *A. thaliana* and *O. sativa*, the impact of duplication events was evaluated on *Brd*-genes among genomes of 79 photosynthetic organisms, including monocots and dicots. *Brd*-gene copies among four lower organisms ranged from 09-16, while *P. abies* harboured 28 copies with no evidence of duplications (Figure 11A). *A. trichopoda* harbored one tandem-duplicate, while *P. patens* showed four block and eight tandem duplicated *Brd*-genes (Figure 11A). Among the monocots, *Brd*-gene copies ranged from 14 (*A. shenzhenica*) to 79 (*T. aestivum*), and except three, all genomes showed different duplication types, a) block events (BD), b) tandem events (TD), c) both tandem and block events (TD, BD), d) tandem and combined events (TD, TD + BD), e) block and combined events (BD, TD + BD), and f) all events (Figure 11A). Events BD, TD and TD + BD were responsible for higher *Brd*-genes in several monocots viz. *Z*. *mays*, *M*. *acuminata*, *E*. *guineensis*, *M*. *sinesis*, *T*. *turgidum*, *S*. *spontaneum* and *T*. *aestivum* (Figure 11B). Likewise, dicots also showed several combinations of events (BD, BD and TD, TD + BD) leading to *Brd*-genes from 20 (*B*. *vulgaris*) to 62 (*G*. *max*)’ Events BD, TD and TD + BD were major contributors to higher *Brd*-genes among *D. carota, P. trichocarpa, M. esculanta, C. arietinum, B. rapa, B. oleracea, C. quinoa, P. bretscheneideri, A. chinensis* and *G. max* (Figure 11C). The duplication events seems to have contributed towards expansion of *Brd*-gene copies among plants.

## 4. Discussion

The modulation of chromatin state by coordinated action of epigenetic mark readers, writers and erasers is central to cellular responses towards metabolic, developmental and environmental cues (Bourbousse et al. 2020; Lauria and Rossi, 2011; Loidl, 2004; Ojolo et al. 2018; Samo et al. 2021; Strahl and Allis, 2000). Studies on Brd-family of epigenetic mark readers, predominantly from animal systems, show their importance in cellular functions important for diverse conditions (Boyson et al. 2021; Tamkun et al. 1992; Rao et al. 2014; Sanchez and Zhou, 2009; Taniguchi, 2016; Uppal et al. 2019; Zeng and Zhou, 2002). On the other hand, plant *Brd*-homologs are relatively less explored, with few studies on certain types viz. GTE (Airoldi et al. 2010; Chua et al. 2005; Duque and Chua, 2003; Misra et al. 2017), GCN5 (Martel, et al. 2017), and SANT type (Sukarta et al. 2020). Recent report has shown three AtBRDs as subunit of SWI/SNF multi-protein chromatic remodeler (Jarończyk et al., 2021). Hence, the understanding significance of diverse Brd-family advocates a thorough analysis. The present comparative analysis of *A. thaliana* and *O. sativa Brd*-homologs provided insights into important aspects viz. diversity of genes/proteins/regulatory elements, orthologs and paralogs, duplication and AS-mediated effects on key Brd features, while expression analysis of duplicate-members corroborated the promoter differences. The duplication events were also found to be important contributors towards *Brd*-family expansion among diverse plants.

Presence of multiple At- and OsBrd-members suggests their involvement in diverse cellular functions (as in humans), however, the number of homologs and domain diversity was substantially less (Filippakopoulos et al. 2012; Fujisawa and Filippakopoulos, 2016), suggesting that certain Brds (and associated functions) may be specific to animals. Further, in both plants, the Brd-members harbored a single BRD (BRD-fold) and lacked dual/poly BRD architecture like human BRDs (Filippakopoulos et al. 2012; Sanchez and Zhou, 2009). Interestingly, one OsBRD (^BD2^OsBRDPG3c) was predicted to harbor additional BRD (overlapped with ET domain), however it showed substantial heterogeneity, and lacked key BRD-fold elements. Few lower photosynthetic organisms do harbor Brd-members with more than one BRD viz. MCO15G409l (*M. commoda*) and Cre05.g247000BRD (*C*. *reinhardtii*).

A notable feature of *A. thaliana* and *O. sativa* Brd-members was enhanced diversity due to duplications, that are known to be important for evolution of multi-member gene families among plants (Barker et al. 2012; Flagel and Wendel, 2009; Qiao et al, 2019). Although, the duplicated Brds showed different divergence levels, in general, the *OsBrd*-duplicates were relatively more divergent in regulatory regions, gene/protein organization, and domains. Different outcomes were observed for the tandem duplication (TD) events affecting OsBrds. While, the TD1 event generated a Brd-copy in a 3-member PG3 group, another event affected the *OsBrdST2* (LOC_Os02g09920, domains: BRD-PHD-WHIM1-ZnF) and generated LOC_Os02g09910 that lacked BRD domain (Supplementary Figure 7). Among the six of the total 13 ortholog groups, both species showed single gene representation of certain BRD types viz. GTE1 TF, GTE12 TF, GCN5, BRM, ATPase-family (Figure 1C). The lineage-specific duplications in different OGs/PGs resulted in increased copies of *Brd*-genes encoding primarily TFs of GTE-type (*A. thaliana*, OG1: GTE8; PG1: GTE3; PG2: GTE2 and *O. sativa*, PG3: GTE7) as well as BRPF3 (*A*. thaliana, OG3) and BDF2 (OG5). Both post-speciation duplication, and post-duplication loss can lead to differences in copy-number of genes among plants (Altenhoff et al., 2019; Qiao et al, 2019), such as possibility affecting *AtBrd* or *OsBrd*) members cannot be ruled out completely. Such events may result into divergence of certain gene-copies affecting regulatory, structural and functional.

Changes in promoter structure, *cis*-elements, and CpG islands (initiates dispersed transcription initiation events, Deaton and Bird, 2011) may affect expression dynamics of genes in diverse conditions, as evident in duplicate *AtBrd* and *OsBrd*-genes. Such expression changes can modulate the relative levels of BRDs and alter chromatin dynamics during reprogramming of cellular machinery in response to metabolic and environmental cues (Chang et al. 2020; Lämke and Bäurle, 2017; Ojolo et al. 2018; Zhang et al. 2018). Intriguingly, several *At*- and *OsBrds* were displayed AS-events that is known to enhance transcriptome and/or proteome diversity (Ali et al. 2007; Syed et al. 2012; Reddy et al. 2013; Laloum et al. 2018). The *Brd*-genes the two species specific to different OGs/PGs showed relatively conserved gene/protein organization, however they evolved differently towards the AS-events. Moreover, in the OGs where *Brds* of both species showed AS, the events showed different impact on transcript and/or protein isoforms, and the same was observed for the duplicated *Brd*-members. While AS-events in the UTRs may affect the stability, translation, localization of transcripts (Mignone et al. 2002), in the coding regions, they can alter structural-functional characteristics. Abundance of AS-isoforms of certain *OsBrds* (^BD1^*OsBrd4a.2*, *OsBrd4b.2*, ^BD3^*OsBrd5a.2*) over constitutive transcripts may have some significance (Supplementary Figure 8), which needs to further investigated for better insights. Previous reports have also indicated that different duplicated genes may display independent, functionally shared, or accelerated AS-modes (Iñiguez, 2017). Both *A. thaliana* and *O. sativa Brd*-duplicates generates non-shared AS-isoforms indicates their divergence towards sub-functionalization (Iñiguez, 2017). Among few studies on plant Brds, AS-mediated modulation of fate and interaction of two similarly localized GCN5 isoforms was reported in *B. distachyon* (Martel et al. 2017). As like duplications, AS-events are also contributing towards enhancing the diversity of Brd-homologs in both the plants, deciphering the significance of ASs in functioning of *AtBrd* and *OsBrd*-homologs is worth investigating.

The BRD/BRD-fold interacts with acetylated lysine on histones, where structural variations can alter this interaction, and associated functions of the Brd-proteins (Josling et al. 2012). Present analysis identified *A. thaliana* and *O. sativa* BRD-members (including duplicates) with variations like substitutions at key conserved sites, additional secondary elements, and partial/complete loss of BRD-fold elements, which might be critical for interaction capability/affinity with the chromatin. However, in view of their tissue-specific expression dynamics, it will be important to decipher their structural-functional characteristics vis-à-vis other Brd-members. Although, a long N-ter deletion that caused loss of αZ-helix, ZA-loop in OsBRD3a and OsBRDST2 was not observed among AtBRDs, an uncharacterized human protein showed similar deletion and loss of elements (Supplementary Figure 6). Interestingly, the At- and OsBRDs BRD-fold elements harboured several conserved signatures, including leucine repeat pattern in αZ and sites in ZA-loop, suggesting similar roles in interaction with αC and loop stabilization, as reported in human-BRDs (Filippakopoulos et al., 2012). However, plant specific site variations in αB, αC and ZA loop, particularly among BRD-duplicates might affect their interaction with chromatin, and associated functions, which will need further investigations.

The consistency between the BRD-region based relationships among At- and OsBRDs and ortholog-paralog clustering, is indicative of its utility in deciphering the divergence of Brd-family in a species, and also to overcome issue related to the analysis of full-length multi-domain proteins (Nakano et al. 2006). The At/OsBRD-homologs lacked dual-BRDs like human BRD2, BRD3, BRD4, TAF1, WDR9 (Filippakopoulos et al. 2012), however, similar domains among different At/OsBRDs were identified, and it would be interesting to find out if they differ in interaction capability as seen for domains of dual-BRD proteins (Miller et al. 2016).

Contribution of genomic duplications in *Brd*-gene copy number was also evident in most plants analyzed in the study, as such events are known to enhance the copy number and/or diversity of plant genes (Qiao et al. 2019). No duplications were evident among lower photosynthetic organisms (*M. commoda*, *S. moellendorffii*, *C. reinhardtii*, *M. polymorpha*) that harbor with few *Brd*-genes. The gene copies increased in *A. trichopoda* (single genome duplication event, Amborella Genome Project, 2013) and *P. patens* with two whole genome duplication events (Lang et al. 2017). Interestingly, without duplications *P. abies* contain higher *Brd*-gene copies, which might be due to inherent transposon activity, also attributed to its large genome size (Nysted et al. 2020). Among higher plants, more duplication events have contributed towards higher gene copies (Qiao et al. 2019). Monocots affected by multiple duplication events (ζ, ancestral; ε, paleohexaploidization; σ and ρ, predating Poaceae divergence), lineage-specific events (*M. acuminata*), and polyploidy (*T. turgidum*, tetraploid; *T. aestivum*, hexaploid) harbor higher *Brd*-gene copies. Likewise, the dicots affected by primitive duplications (ζ, ε), triplication (WGT, γ), and lineage-specific WGD/ploidy events (α and β, crucifer lineage; *Gossypium*-specific ploidy; WGDs specific to poplar, legumes, *Glycine*) also showed higher *Brd*-gene copies. Moreover, *Brd*-gene copies may also be affected by post-duplication losses/deletions (Qiao et al. 2019), and is likely in plants like *A. shenzhenica*, *P. equestris*, *Z. marina*, which lack *Brd*-duplicates despite an ancient WGD event (Cai et al. 2014; Olsen et al. 2016; Zhang et al. 2017).

Overall, the present analysis revealed extensive diversity among the *A. thaliana* and *O. sativa Brd*-members w.r.t. gene/proteins, promoters, domains/motifs, expression pattern, and sequence/structural differences in the BRD-fold. Analysis identified functionally conserved orthologs, species-specific evolutionary trend, and involvement of duplication and AS-events towards enhanced diversity of *Brd*-homologs. The duplication-mediated *Brd*-gene copies were substantially enhanced among complex photosynthetic organisms with history of duplication events. The plant *Brd*-gene family is relatively less studied, however its diversity, impact of duplication and AS-events, domain signatures, suggests involvement in diverse cellular mechanisms, and needs a thorough analysis for understanding their functional significance.

## Authors contribution

ATV: analysis of gene and protein sequences, RNA-Seq data, phylogeny, RPS: analysis of splicing events and domain heterogeneity; HSM: data analysis and review, manuscript writing; AS: planning and execution, data analysis and review, manuscript compilation and communication.

## Data availability

The sequences used in the study are available from the public databases, GenBank-NCBI (https://www.ncbi.nlm.nih.gov/genbank), Rice Genome Annotation Project (RGAP, http://rice.uga.edu/), The Arabidopsis Information Resource (TAIR, https://www.arabidopsis.org/index.jsp) and PLAZA monocot and dicots online resources (https://bioinformatics.psb.ugent.be/plaza/). All the locus numbers and accession numbers are listed in manuscript, Table 1 and Table 2.

## Supplementary data files

Supplementary Table 1: Oligonucleotide primers used for RT-qPCR analysis of *AtBrd* and *OsBrd* genes.

Supplementary Table 2: List of conserved motifs identified among 28 *A. thaliana* BRD-proteins. Supplementary Table 3: List of conserved motifs identified among 22 *O. sativa* BRD-proteins.

Supplementary Figure 1: Schematic representation of alternative splicing events affecting BRD-genes of *A. thaliana* (A) and *O. sativa* (B).

Supplementary Figure 2: Schematic representation of diversity of motifs among BRD-proteins of of *A. thaliana* (A) and *O. sativa* (B).

Supplementary Figure 3: Diversity of *cis*-elements in the upstream regions of *A. thaliana Brd*-genes as per analysis as PlantCARE database.

Supplementary Figure 4: Diversity of *cis*-elements in the upstream regions of *Oryza sativa Brd*-genes as per analysis as PlantCARE database.

Supplementary Figure 5: Multiple sequence alignment of the bromodomain-regions of 28 AtBRDs and 22 OsBRD homologs.

Supplementary Figure 6: Comparison of homology model of a normal BRD-region with typical features of the BRD-fold and homology model of BRD-region of a human protein (K2026_Human, UniProt ID: Q5HYC2) with a deletion at N-terminal region leading to loss of αZ and ZA-loop elements.

Supplementary Figure 7: CDD-NCBI based conserved domain analysis of OsBrd4 (LOC_Os02g09920, BRD-homolog with BRD-PHD combination and its tandem duplicate (LOC_Os02g09910) encoding protein with only PHD.

Supplementary Figure 8: Heatmap-based analysis of RNA-Seq data of constitutive and alternative transcripts of *OsBrd*-genes in different tissues and few stress conditions.

## Acknowledgements

Authors thank Dr Sheetal Uppal, Molecular Biology Division, Bhabha Atomic Research Centre for suggestions and comments.

