## Supplementary Table 1 for "Genome-wide Analysis of Bromodomain Gene Family in Arabidopsis and Rice"

Supplementary Table 1: Oligonucleotide primers used for RT-qPCR analysis of *AtBrd* and *OsBrd* genes.

| S. No. | Primer Name | Gene | Primer Sequence (5´-3´) |
|---|---|---|---|
| 1 | AtBrd3-ForP | *AtBrd3* | AGTTACTGAATCAGGAAAAGC |
| 2 | AtBrd3-RevP | (AT1G17790) | GTCACTGTCAGTATCACTAGAG |
| 3 | AtBrd8-ForP | *AtBrd8* | GTCATGTGGCATCTACTGTTC |
| 4 | AtBrd8-RevP | (AT1G73150) | AGCTATCACTGTCAGAGCCACTA |
| 5 | AtBrd11-ForP | *AtBrd11* | GACTCGAACGTCAGGAAACAATTG |
| 6 | AtBrd11-RevP | (AT2G42150) | AACTCTGAAGATCCTCTGTG |
| 7 | AtBrd12-ForP | *AtBrd12* | AGGGTGGTGCCGGCGAGATC |
| 8 | AtBrd12-RevP | (AT2G44430) | AATCCTTAGTGTCCTGGCTC |
| 9 | AtBrd20-ForP | *AtBrd20* | GTAACCTGGTCTCGCCAGGATC |
| 10 | AtBrd20-RevP | (AT3G57980) | TTCAACTGCAATGACGAGATGG |
| 11 | AtBrd21-ForP | *AtBrd21* | AGAGGAAAGGACAGAAATAC |
| 12 | AtBrd21-RevP | (AT3G60110) | AATCCTTAGTGTCCTGGCTC |
| 13 | AtBrd22-ForP | *AtBrd22* | CTCCACCAAGAAACATGCCTCCGG |
| 14 | AtBrd22-RevP | (AT5G10550) | TAGAGCCACCACTAGCAGCGGCAGCTG |
| 15 | AtBrd28-ForP | *AtBrd28* | CTCCACCTAGGAACATGGCTTC |
| 16 | AtBrd28-RevP | (AT5G65630) | CACTACTAGCAGCAGCTGCAA |
| 17 | AtAct-ForP | *AtActin* | CACAGCACTTGCACCAAGCAGCAT |
| 18 | AtAct-RevP | (AT3G18780) | TGGAGATCCACATCTGCTGGAATG |
| 19 | OsBrd1-ForP | *OsBrd1* | ATGATAATGAGAGGTATGTAGGCTCATCATCACC |
| 20 | OsBrd1-RevP | (LOC_Os01g11580) | CACTAGACGAAGAGCCTGAATCGC |
| 21 | OsBrd2-ForP | *OsBrd2* | CGGGATGATGGAGGTGGACCTGGA |
| 22 | OsBrd2-RevP | (LOC_Os01g46040) | TCATGGCTCCTGCTTCACCTTAACC |
| 23 | OsBrd10-ForP | *OsBrd10* | CCGCTACGATGGTGGACAACGGTGATGTGACG |
| 24 | OsBrd10-RevP | (LOC_Os04g53130) | GGGCGGACCCAGGGCCAGGACA |
| 25 | OsBrd11-ForP | *OsBrd11* | ACAATGGCAATAGAGAGCAAGGATCCTGA |
| 26 | OsBrd11-RevP | (LOC_Os04g53170) | CCTCAGTATCCTTCTCAATCTCCACTGATTGGT |
| 27 | OsBrd13-ForP | *OsBrd13* | AGAGCTTATGAAACGGGTGAAGAAGGGCTTCA |
| 28 | OsBrd13-RevP | (LOC_Os06g24870) | TCATCGCCTGTGTTATCATTGCCACCATTCTCCTG |
| 29 | OsBrd17-ForP | *OsBrd17* | GAGGATGAAGGCTCCAGACCATACTTTGACG |
| 30 | OsBrd17-RevP | (LOC_Os08g01794) | TGGCTTGTTTCTGTTGGCTCCTTATGTTGG |
| 31 | OsBrd18-ForP | *OsBrd18* | GATATTGGCGATGAGATGCCGACGGCA |
| 32 | OsBrd18-RevP | (LOC_Os08g09340) | GAGCTCCTCGAGTCAGAATCACTGGATG |
| 31 | OsBrd19-ForP | *OsBrd19* | GAAATACGATCTCTCGATCGGGTCGTTGCAATCA |
| 32 | OsBrd29-RevP | (LOC_Os08g39980) | CGCCGGCCTCCTCCGGTGATCC |
| 33 | OsBrd20-ForP | *OsBrd20* | CGCCTGTTGCTCAGCTGGTATCTG |
| 34 | OsBrd20-RevP | (LOC_Os09g33980) | TGGTGGATTGTCAGTTTCTTTCAGAACACC |
| 35 | OselF1α-ForP | *OselF1α* | GCTGCAACAAGATGGATGCCACCA |
| 36 | OselF1α-RevP | (LOC_Os03g08010) | GAAGGGAATCTTGTCAGGGTTGTAGC |

Note: The locus numbers of the *AtBrd* genes are as per The Arabidopsis Information Resource (TAIR, https://www.arabidopsis.org/) and of *OsBrd* genes, as per Rice Genome Annotation Project (RGAP, http://rice.uga.edu/).
