## Supplementary Table 2 for "Genome-wide Analysis of Bromodomain Gene Family in Arabidopsis and Rice"

Supplementary Table 2: List of conserved motifs identified among twenty-eight *Arabidopsis thaliana* bromodomain (BRD) containing proteins at per analysis using MEME suite software.

| Motif | Motif Sequence | E-Value | Sites | Width | Functional Annotation |
|---|---|---|---|---|---|
| M1 | EJPDYYNIIKHPMDLGTIKKKLEKG | 3.10E-213 | 24 | 25 | BRD |
| M2 | YSSPLEFAADVRLTFNNAMTY | 1.10E-166 | 24 | 21 | BRD |
| M3 | MKQCETLLRKLMKHKHGWVFNTPVDVVGL | 1.90E-134 | 12 | 29 | BRD |
| M4 | NPEGNDVYVMAEKLLKLFEERWKTIEKKY | 3.80E-87 | 12 | 29 | BRD |
| M5 | QIVKKSNPELSQGDDEIEJDIDALDDETLWELRRF VDEYLK | 9.30E-83 | 12 | 41 | ET |
| M6 | QTWGTWEELLLACAVKRHGTGDWDSVASEVQ | 2.80E-47 | 5 | 31 | Myb-like DBD |
| M7 | EPAKRDMTDEEKRKLGEDLQSLPPDKLZ | 2.10E-33 | 11 | 28 | ET |
| M8 | PWLEELRKLRVAELRREVERYDLSINSLQLKVKK LEEERE | 3.90E-32 | 7 | 40 | - |
| M9 | AAEAKRKRELEREAARQALLEMEKSVEINENSRF LEDLELL | 2.20E-29 | 4 | 41 | - |
| M10 | DGVSKMVLSLSGLSSSERRELKRRLKSELEZVRSL RKRIE | 9.40E-28 | 8 | 40 | BRD |
| M11 | PEKRYRAAJLKNRFADIILKAREKPLNQN | 5.70E-25 | 5 | 29 | - |
| M12 | PVGLNAEYGYARSLARYAANLGPVAWKIASQRI EKVLPSGIKFGRGWVGE | 4.30E-27 | 3 | 50 | - |
| M13 | KGDPEKLQREREELELQKKKEKARLQAEAKAAE EARRKA | 1.60E-24 | 5 | 39 | - |
| M14 | ICVWNAADGSLVHCLTGHSESSYVLDVHPFNPRI AMSAGYDGKTIIWDIW | 2.10E-21 | 2 | 50 | WD |
| M15 | WFLHDNFVTCSRDGRANIWRPEIRGVGGPSGRGL GFYHLKVPPPPLPP | 4.10E-29 | 4 | 48 | TFIID, subunit TAF1 |

Analysis of conserved motifs was carried out online using MEME software, available at MEME online Suite (version 5.4.1, http://meme-suite.org/tools/meme,Timothy et al., 2015).
