## Supplementary Table 3 for "Genome-wide Analysis of Bromodomain Gene Family in Arabidopsis and Rice"

Supplementary Table 3: List of conserved motifs identified among twenty-two *Oryza sativa* bromodomain (BRD) containing proteins, as per analysis using MEME suite software.

| Motif | Motif Sequence | E-Value | Sites | Width | Functional Annotation |
|---|---|---|---|---|---|
| M1 | YGSLEEFAADVRLTFSNAMTYNPKGHDVH | 9.10E-187 | 22 | 29 | BRD |
| M2 | DVVALGJPDYFDIIKKPMDLGTIRKKLE | 2.00E-145 | 15 | 28 | BRD |
| M3 | LPPEKLDNVLQIVKKRNGSPELVGDEIELDIDEMD VETLWELDRFVANYK | 6.80E-84 | 7 | 50 | ET |
| M4 | LKRCGEILKKLMKHKAAEPFNTPV | 2.30E-27 | 9 | 24 | BRD |
| M5 | ATFENMYKPAHSWFEQEPKILEPPMPVPPPPEPEK PAPSTV | 4.40E-23 | 5 | 41 | - |
| M6 | MKKPKAREPNKREMTLEEKNKLRVGLZ | 1.10E-20 | 5 | 27 | - |
| M7 | YVDIGDEMPTATYQSVEIEKDTEAASSGSSSSSDS GSSKDSVSESGNAH | 3.90E-15 | 3 | 49 | - |
| M8 | IVNGENADVIDASVANDSDMLVNGSTATMVDNG DVTMAIESKDPDKITTQ | 5.10E-13 | 2 | 50 | - |
| M9 | MKRVKKGFMKNWLAAGLYSDVQENGGNDNTG DEDVKGSKGKSKQKRRRLG | 4.00E-11 | 2 | 50 | - |
| M10 | MAEQLLEIFEAKWPEIEAKV | 6.10E-07 | 6 | 20 | BRD |
| M11 | VQNAKPKVYSRVRLKFKSAKVLETHQGPSEAKA PVDGGGGKPASAAPEA | 5.70E-06 | 2 | 49 | - |
| M12 | HHHHHHQW | 9.00E-06 | 3 | 8 | - |
| M13 | NQDRKADSVSEPLPSKQETVLENVESETALEPRSS QELEVKQATPERQRD | 5.00E-04 | 2 | 50 | - |
| M14 | VAEKAIVSPDGQKDAQAAELSGSDKDKMARKVA SIKIKSVGLSSVEDK | 4.90E-02 | 2 | 48 | - |
| M15 | MKRKRGRKMGKKGKLKASITADASPMSPSSPST VDASSKSP | 9.80E-01 | 2 | 41 | - |

Analysis of conserved motifs was carried out online using MEME software, available at MEME online Suite (version 5.4.1, http://meme-suite.org/tools/meme,Timothy et al., 2015).
