## Supplementary Figure 1 for "Genome-wide Analysis of Bromodomain Gene Family in Arabidopsis and Rice"

Supplementary Figure 1: Schematic representation of alternative splicing events in *Brd*-genes of *A. thaliana* (A) and *O. sativa* (B), belonging to different ortholog groups (OGs), paralog groups (PGs), and singleton category (STs). Block duplicated genes (BD), constitutive transcript (.1), alternative transcripts (.2 to .6), UTRs (white boxes), exons (dark grey boxes) and introns (dashed lines) are indicated in the figure. Scale on the top indicates the length (kilobase, kb).

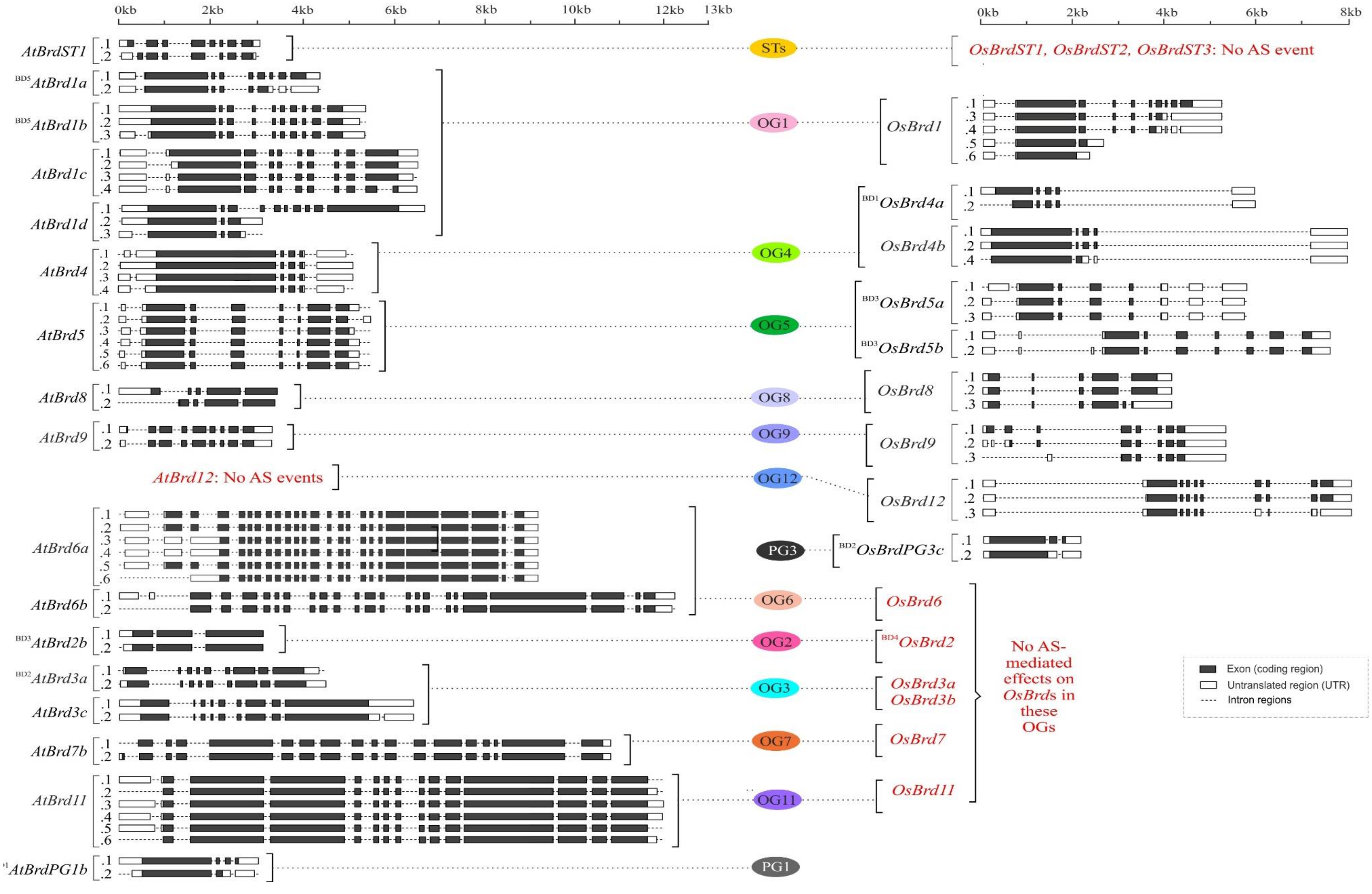


(A) (B)
