## Supplementary Figure 2 for "Genome-wide Analysis of Bromodomain Gene Family in Arabidopsis and Rice"

Supplementary Figure 2: Motif heterogeneity among Brd-proteins of *A. thaliana* (A) and *O. sativa* (B), belonging to thirteen ortholog groups (OG1-13), three paralog groups (PG1-3), and singleton category (STs). Motifs M1-M15 are shown in different color coded. Scale on the top indicates the protein length (number of amino acids). Duplicate BRD-pairs are indicated with the designations 'BD' (block duplication) and 'TD' (tandem duplication) in the names.

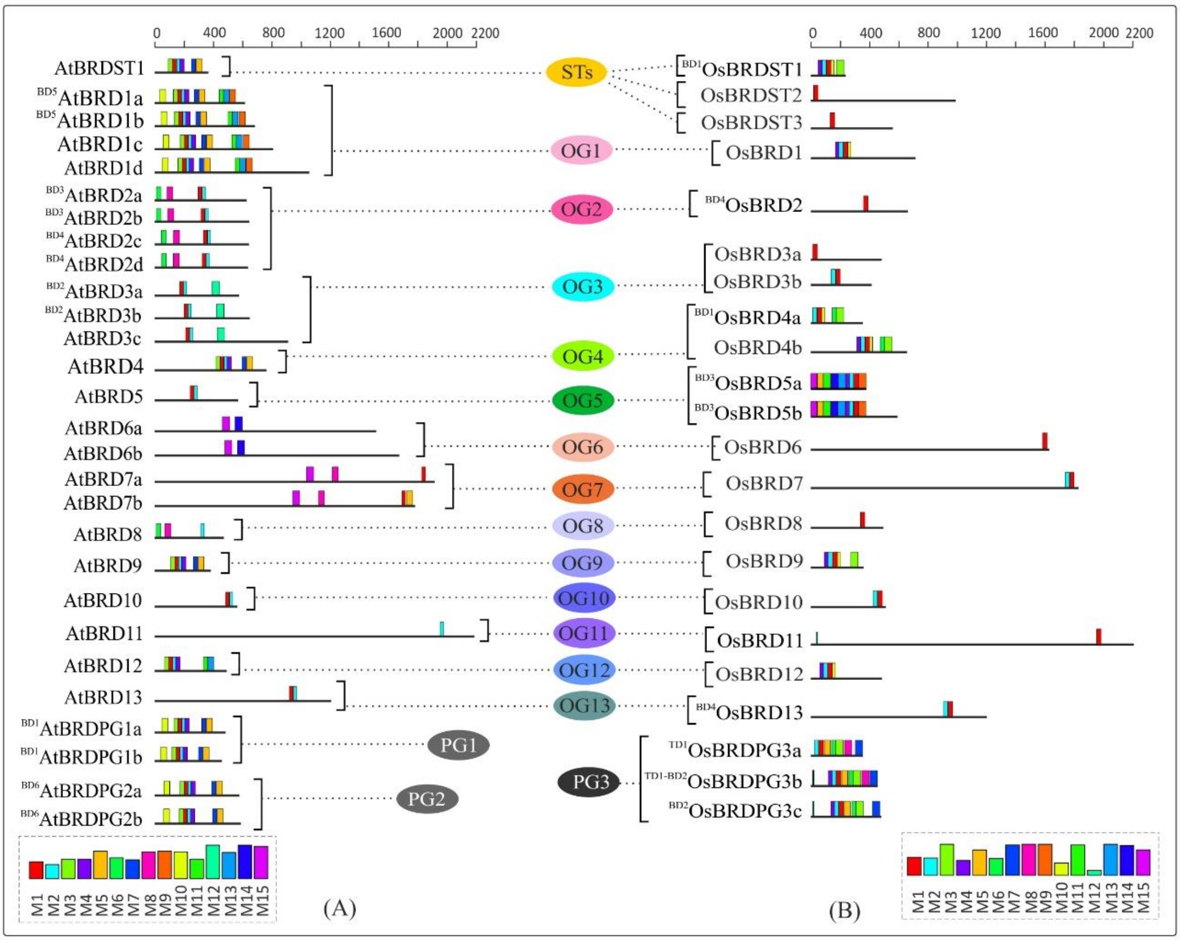
