## Supplementary Figure 3 for "Genome-wide Analysis of Bromodomain Gene Family in Arabidopsis and Rice"

Supplementary Figure 3: Diversity of *cis*-elements (as per analysis at PlantCARE database) in the upstream regions of *A. thaliana Brd*-genes belonging to thirteen ortholog groups (OG1-13), two paralog groups (PG1-2), and singleton category (STs). Different types of elements are indicated by different symbols/colours, *cis*-elements belonging to six major categories are indicated below, and block-duplicated genes are indicated by the designation ‘BD’ in the gene name. Scale on the top indicates the length in kilobase.

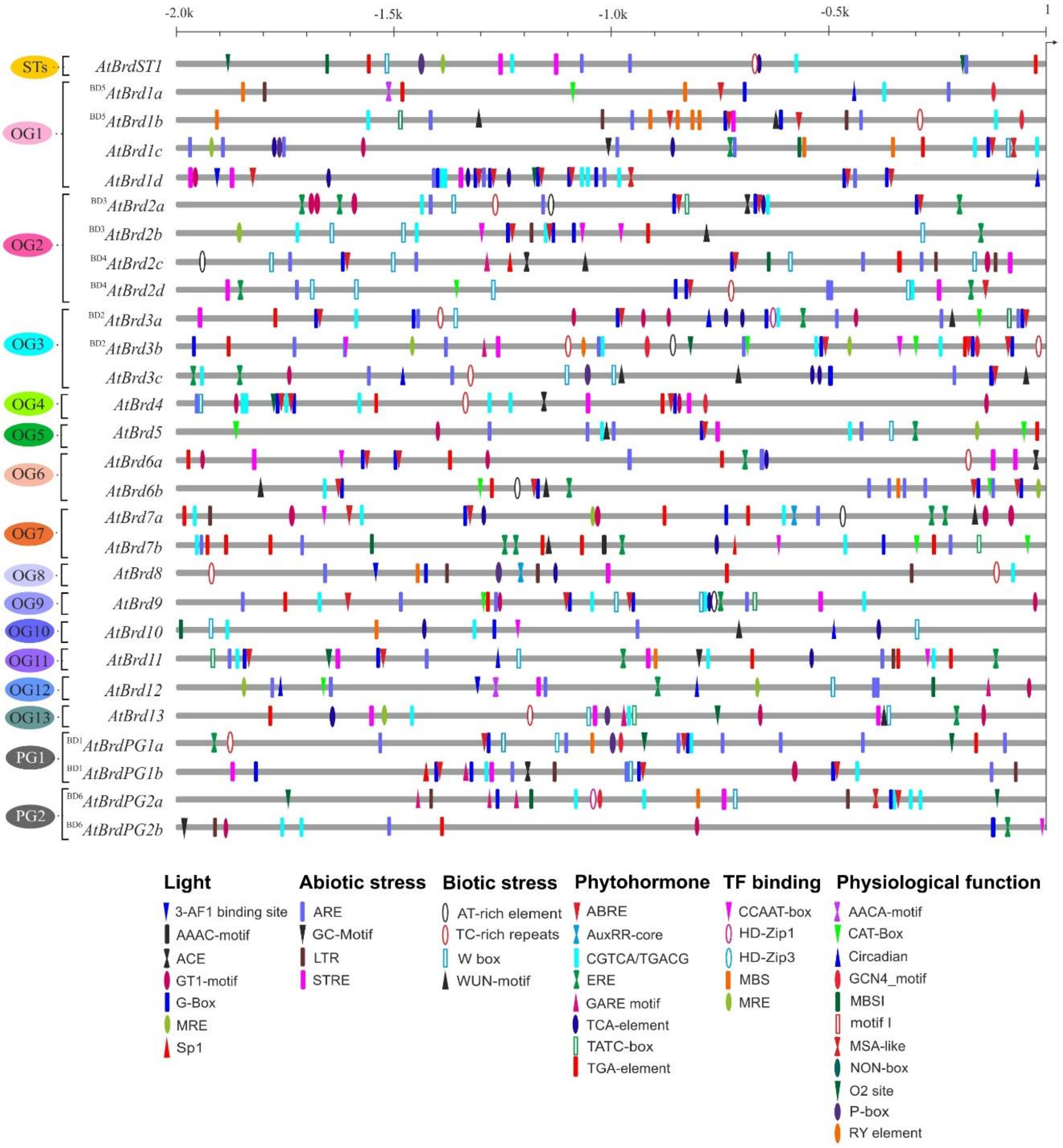
