## Supplementary Figure 5 for "Genome-wide Analysis of Bromodomain Gene Family in Arabidopsis and Rice"

Supplementary Figure 4: Diversity of *cis*-elements in the upstream regions of *O. sativa Brd*-genes belonging to thirteen ortholog groups (OG1-13), one paralog group (PG3), and singleton category (STs), as per analysis at PlantCARE database. Different types of elements are indicated by different symbols/colours, *cis*-elements specific to six functional categories are listed below, and genes affected by block or tandem duplications are indicated by the designation ‘BD’ or ‘TD’ in the gene names. Scale on the top indicates the length in kilobase.

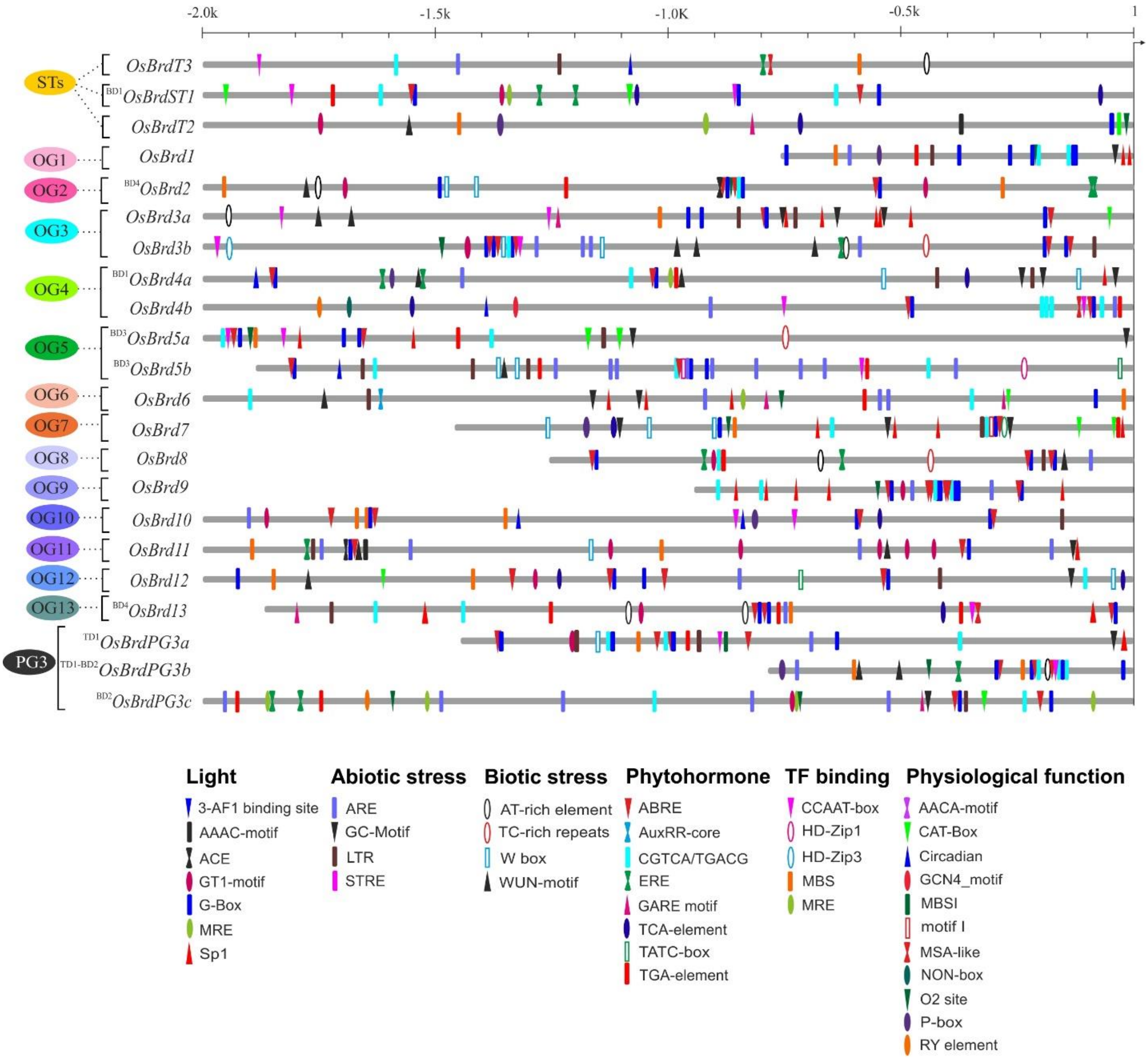
