## Supplementary Figure 5 for "Genome-wide Analysis of Bromodomain Gene Family in Arabidopsis and Rice"

Supplementary Figure 5: Multiple sequence alignment of the bromodomain-region of 28 AtBRD and 22 OsBRD-homologs, as per details in the Table 1 and Table 2. Scale on the top indicates the sites in the MSA file. The duplicate-pairs are indicated with names with designations ‘BD’ (block duplication) and ‘TD’ (tandem duplication), while numerals (I-VI) indicates the six major clusters as per phylogenetic analysis (Figure 9A).

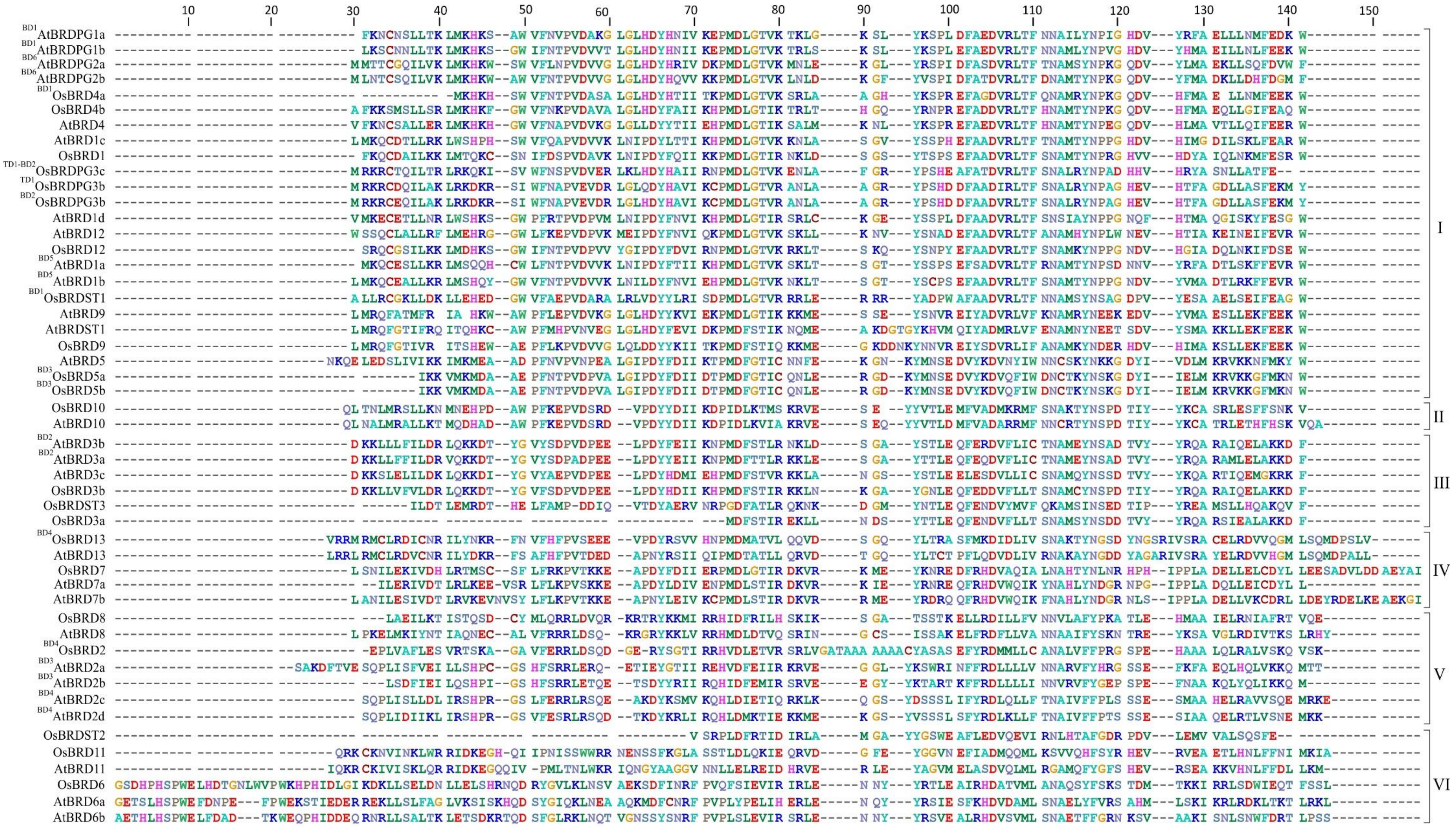
