## Supplementary Figure 6 for "Genome-wide Analysis of Bromodomain Gene Family in Arabidopsis and Rice"

Supplementary Figure 6: (A) Homology model of a normal bromodomain (BRD) region containing all typical structural features of the BRD-fold (four α helices: αZ, αA, αB, αB and three loops: ZA, AB, BC). (B) Homology model of BRD-region of a human protein (K2026_Human, UniProt ID: Q5HYC2) with a long deletion at N-terminal region (similar to deletion in OsBRD4 and OsBRD7), leading to loss of αZ and ZA-loop elements.

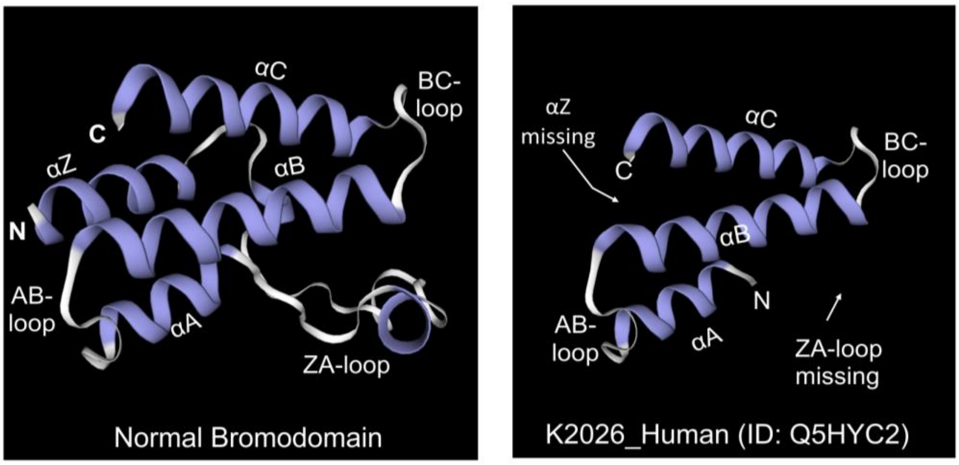


(A) (B)
