## Supplementary Figure 7 for "Genome-wide Analysis of Bromodomain Gene Family in Arabidopsis and Rice"

Supplementary Figure 7: CDD-NCBI based conserved domain analysis of OsBRDST2 (LOC_Os02g09920, BRD-homolog with BRD-PHD-WHIM1-ZnF domain combination (A) and its tandem duplicate (LOC_Os02g09910) encoding protein contaning only PHD domain (B).

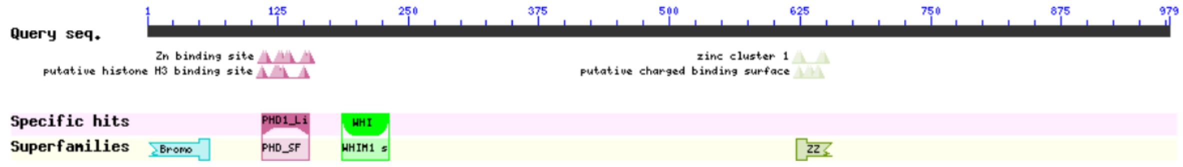


(A)

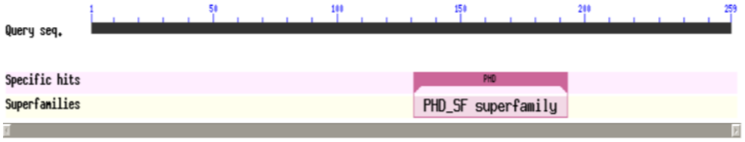


(B)
