## Supplementary Figure 8 for "Genome-wide Analysis of Bromodomain Gene Family in Arabidopsis and Rice"

Supplementary Figure 8: Heatmap-based analysis of RNA-Seq data of constitutive and alternative transcripts of *OsBrd*-genes in different tissues (A) and two stress conditions (cadmium and drought, DS) (B). The *Brd* transcript designations (constitutive transcript: .1; alternative transcripts: .2 to .5) are mentioned on the left side, tissues and stress conditions are listed on the top, and a gradient color scale indicating expression level from blue (low) to red (high) is shown on the bottom of the figure. Red boex highlight the genes where, in general, level of alternative transcript (0.2) is higher than the constitutive transcript (.1).

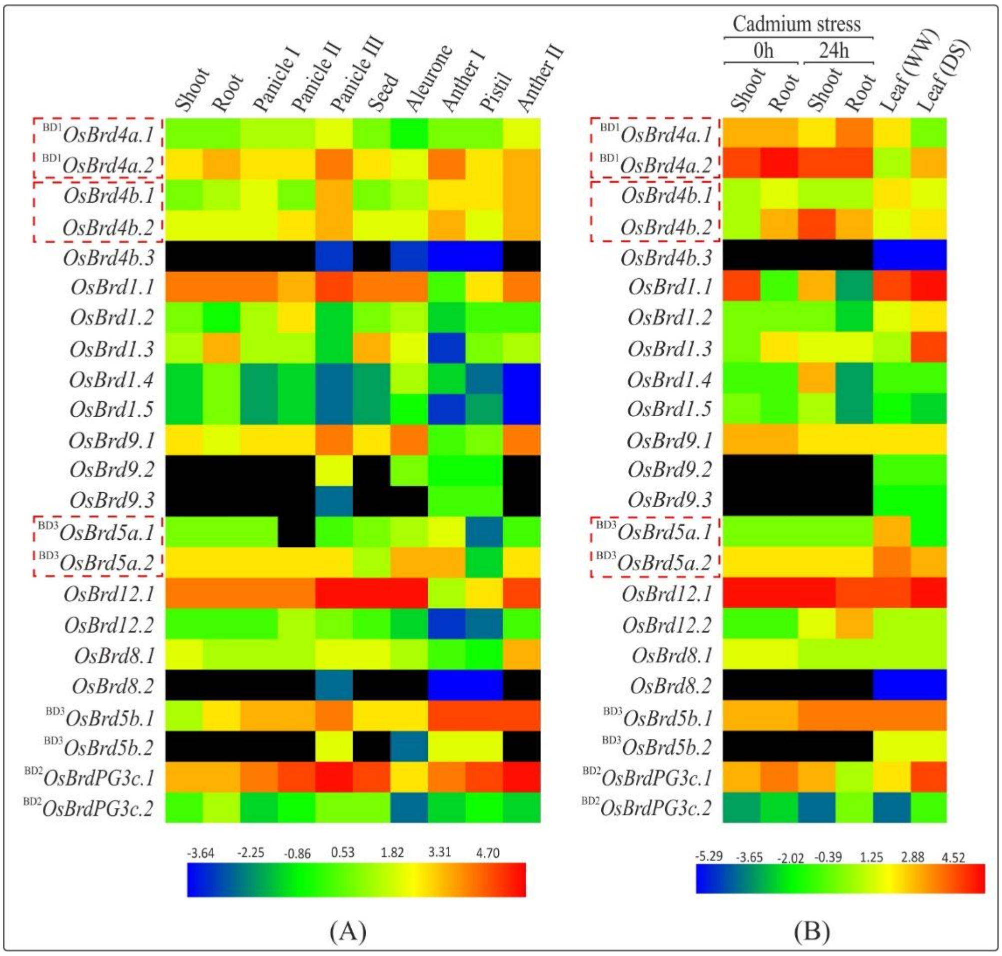
